# Integrated Multi-Omics Enabled by Sequential Extraction for Comprehensive Molecular Profiling of Small Extracellular Vesicles

**DOI:** 10.64898/2026.02.19.706688

**Authors:** Andrew J. Perciaccante, Holden T. Rogers, Yanlong Zhu, Aditi Barnwal, David Inman, Man-Di Wang, Song Jin, Suzanne M. Ponik, Ying Ge

## Abstract

Small extracellular vesicles (sEVs) are membrane-bound particles whose protein, lipid, and metabolite cargo reflects the molecular state of their cells of origin, making them attractive targets for biomarker discovery and therapeutic development. However, comprehensive characterization of sEVs remains challenging due to the extremely limited material available. Here, we present an integrated mass spectrometry-based multi-omics platform for simultaneous characterization of lipids, metabolites, and proteins from a single sEV sample enabled by sequential extraction, maximizing sample utilization. To enhance molecular coverage and analytical depth, the platform combines iterative tandem mass spectrometry for improved small-molecule fragmentation and nano-flow proteomics with data-independent acquisition. We achieved deep and reproducible multi-omic characterization of proteins, lipids, and metabolites using 10 million sEVs. We further demonstrated the compatibility of our multi-omics platform with sEVs isolated from plasma by ultracentrifugation, size-exclusion chromatography with ultrafiltration, and polymer precipitation, revealing purification-dependent differences in molecular profiles associated with tradeoffs in yield and purity of sEVs. By enabling integrated multi-omics from the same sample, this strategy addresses a key challenge in low-input sEV analysis and establishes a robust analytical foundation for synergistic biomarker discovery and therapeutic applications.

## Introduction

Extracellular vesicles (EVs) are membrane-encased structures secreted into the extracellular space by all cell types.^1–5^ Small extracellular vesicles (sEVs) are a subset of EVs that are less than 200 nm in diameter, and include both ectosomes and exosomes.^1–6^ In contrast to ectosomes, which are formed by the budding off of the plasma membrane, exosomes are generated through the endosomal pathway within multivesicular bodies.^3–7^ sEVs play central roles in intercellular communication, and are involved in various pathophysiological processes, including in disease pathogenesis and immune response.^1,6–13^ In cancer, sEVs are recognized for their diverse functions, including promoting angiogenesis, metastasis, and therapeutic resistance.^1,4,6,9–13^ Moreover, cancer cells have been shown to secrete more sEVs than healthy cells, and circulating sEV concentrations in cancer patients can be significantly elevated relative to healthy individuals.^6,9,11,14^ As sEVs carry molecular cargoes, including proteins, lipids, and metabolites, that reflect both their function and cell of origin,^4,6,11,15^ comprehensive characterization of sEV cargoes offers a powerful avenue for understanding the disease mechanisms and enabling precision diagnostics.

sEVs are present in nearly all biofluids and carry disease-derived molecular information, making them highly attractive targets for liquid biopsy applications ranging from biomarker discovery to therapeutics development.^3,4,6,16^ Blood is the most commonly studied biofluid for studying EVs,^3,15,17^ and sEVs are commonly purified from blood plasma using techniques such as ultracentrifugation (UC), size exclusion chromatography with ultrafiltration (SECUF), and polymer precipitation (PPT); however, these purification approaches rely on distinct separation principles and thus produce sEV populations that differ both in yield and purity, ultimately affecting downstream analyses.^3,5,17–21^

Mass spectrometry (MS)-based multi-omics integrating proteomics, lipidomics, and metabolomics provides comprehensive characterization of sEVs molecular cargoes.^22,23^ This approach provides deeper biological insight by integrating multi-omics data to identify relationships between individual analytes from different -omics layers, as well as enriched and/or perturbed pathways.^24–29^ Furthermore, it enables the development of multi-component biomarker panels with better predictive values compared to proteins alone.^30^ Despite these advantages, multi-omics analysis of sEVs remains technically challenging due to the extremely limited material available. With an average radius more than 100-fold smaller than a MDA-MB-231 (MDA) cell, 10 million sEVs have a similar total biological volume comparable to only a few cells.^31^ Conventional multi-omics strategies frequently rely on separate samples for proteomic, lipidomic, and metabolomic analyses, which can compromise cross-omic biological linkage and introduce technical variability.^24,32,33^ To address these challenges, a unified and sensitive multi-omics platform capable of extracting maximal molecular information from the same sEV sample is urgently needed.

Herein, we develop a multi-omics platform designed for comprehensive molecular profiling of sEVs from the same sample, achieving both broad molecular coverage of proteins, lipids, and metabolites with robust quantitation essential for biomarker discovery. Our method utilizes sequential extraction of lipids, metabolites, and proteins from a single sample, maximizing sample utilization. We first demonstrated sensitive and reproducible multi-omic characterization using 10 million sEVs isolated from MDA cells.

We then evaluated its clinical applicability by comparing plasma-derived sEVs isolated using UC, SECUF, and PPT. Our findings reveal distinct molecular signatures for each isolation technique, reflecting marked differences in sEV yield and purity. Together, this work establishes a unified low-input multi-omics strategy that enables simultaneous characterization of proteins, lipids, and metabolites from a single sample of sEVs, providing a foundation for systems-level understanding of sEV biology and biomarker discovery using limited clinical samples.

### Experimental Section

Additional details regarding the purification of sEVs, specific mass spectrometer parameters, and data processing have been included in the *Supporting Information*.

### Chemicals and Reagents

All reagents were purchased from Millipore Sigma (Burlington, MA), unless otherwise noted. Extraction solvents and mobile phases were prepared with LC-MS grade MilliQ filtered water (Millipore Sigma, Burlington, MA). Methanol (MeOH), acetonitrile (ACN), and isopropanol (IPA) were purchased from Fisher Scientific (Waltham, Massachusetts).

### Purification of sEVs from MDA cells

MDA cells expressing mScarlet-CD63 were cultured in Opti-MEM to minimize contamination from bovine serum EVs for 48 hours. The conditioned media were collected, and cells, cell debris, and microvesicles were removed via centrifugation at 300 × g for 10 minutes, 2,000 × g for 20 minutes, and 10,000 × g for 30 minutes, respectively. The media were then concentrated using a Centricon® Plus-70 device, and sEVs were purified via two rounds of ultracentrifugation (100,000 × g for 4 hours at 4 °C). The sEV pellet was resuspended in DPBS, and the yield and size distribution of particles were measured using nanoparticle tracking analysis (NTA). Ten million particles were aliquoted for multi-omics analysis for each sample.

### Purification of sEVs from Plasma

Plasma was centrifuged at 2,000 *× g* at 4 °C for 20 minutes to remove cells and cell debris. The supernatant was aliquoted into 1.5 mL tubes (200 µL/sample) for purification using PPT, SECUF, and UC.

PPT was performed using the Total Exosome Isolation (from plasma) kit (Invitrogen, Waltham, MA). Aliquoted partially clarified plasma samples were centrifuged at 18,000 *× g* at 4 °C for 30 minutes to remove microvesicles. The supernatants were transferred to new 2.0 mL tubes, and sEV purification was performed according manufacturer instructions, without Proteinase K treatment. Purified sEVs were resuspended in 100 µL of DPBS and snap frozen for later characterization using NTA and multi-omics analysis.

For SECUF purification, aliquoted partially clarified samples were centrifuged at 18,000 *× g* at 4 °C for 30 minutes to remove microvesicles. The supernatants were transferred to new 2.0 mL tubes, and size exclusion chromatography was performed by passing the clarified plasma through qEVoriginal 35 nm Gen 2 SEC columns. Details on column equilibration and cleaning are included in the *Supporting Information*. Purified sEVs were concentrated to less than 200 µL using kDa molecular weight cut-off filters (REF #UFC903008, Millipore, Burlington, MA). The concentrated samples were transferred to clean 2.0 mL tubes, snap frozen, and stored at -80 °C for NTA and multi-omics analysis.

UC samples were prepared by first centrifuging the aliquoted partially clarified plasma at 18,000 *× g* for 30 minutes at 4 °C to remove microvesicles. Supernatants were transferred to ultracentrifuge tubes (344057, Beckman Coulter, Brea, CA), diluted to 3 mL with DPBS, and centrifuged twice at 100,000 *× g* at 4 °C for 2 hours. The pellets were resuspended in 100 µL of DPBS prior to being snap frozen before characterization using NTA and multi-omics analysis.

### Nanoparticle Tracking Analysis

Nanoparticle tracking analysis was performed using the Malvern Panalytical NanoSight NS300 (Malvern, United Kingdom). Isolated sEVs were suspended in one milliliter of Dulbecco’s phosphate buffered saline (DPBS). sEVs isolated using PPT were diluted 20-fold with DPBS to adjust to the working range of the instrument. The concentrations presented herein account for dilution factors.

### Extractions for Multi-omics

A modified Matyash^34^ extraction was performed to simultaneously extract lipids and metabolites while precipitating proteins. Purified sEVs were thawed on ice, and sample volumes were adjusted to 200 µL with chilled water. To each sample, 387 µL of chilled MeOH, containing 4.17 ng acylcarnitine 18:1-d3 (Caymen Chemical, Ann Arbor, MI) and 4.69 ng of each of C12:0-d23, C14:0-d27, C16:1-d14, C18:1-d17, and C18:0-d35, was added. Samples were vortexed, followed by the addition of 1,290 µL of chilled methyl *tert*-butyl ether containing 0.375 µL of Splash Lipidomix (Avanti Polar Lipids, Alabaster, AL), and vortexed again. To induce phase separation, 123 µL of chilled water, containing 0.1 µL of 500 µM NSK-A heavy amino acid mix (Cambridge Isotope, Tewksbury, MA) was added. Samples were vortexed intermittently for ten minutes to maximize extraction efficiency,^35,36^ and kept on ice between vortexing cycles. Samples were centrifuged at 18,000 *× g* at 4 °C for ten minutes. Following phase separation, 1,300 µL of the upper organic layer and 350 µL of the lower aqueous layer were extracted to new tubes and dried under vacuum centrifugation. The organic extracts were resuspended with 1:1 chloroform:MeOH and were stored at -80 °C with the dried aqueous extracts until analysis.The protein pellet was dried under vacuum centrifugation, snap frozen, and stored at -80 °C until further processing.

The protein pellet was resuspended with 24 µL of lysis buffer, containing 5% (w/v) of our in-house synthesized MS-compatible surfactant, Azo,^37–39^ in place of SDS, and 50 mM triethylammonium bicarbonate (TEAB) at pH 8.5. The resuspended pellets were vortexed briefly and bath sonicated for 10 minutes. Proteins were reduced via the addition of 1 µL of 120 mM Tris (2-carboxyethyl) phosphine (TCEP) and incubated at 55 °C for 15 minutes on a thermoshaker operated at 800 RPM. Alkylation was performed using 3.29 µL of 400 mM chloroacetamide, and samples were incubated at room temperature for 30 minutes. Acidification and S-Trap binding, washing, digestion, and elution were performed using S-Traps (Protifi, Fairport, NY) according to the manufacturer’s instructions.^40^ For PPT samples, 10 µg of protein was loaded on the S-Trap, and for all other samples, the entire sample volume was used. Each sample was digested overnight via the addition of 1 µg of Trypsin Gold (Promega, Madison, WI). Following peptide elution, samples were dried using vacuum centrifugation.

### Proteomics

Dried peptides were resuspended with 11 µL of 0.1% formic acid. Peptide concentration was estimated using a NanoDrop One Microvolume UV-vis Spectrophotometer (Thermo Fisher Scientific, Waltham, MA). Using a nanoElute 2 (Bruker, Billerica, MA) nano-flow LC system, a total of 250 ng of peptide was injected onto an Aurora Ultimate C18 column (25 cm x 75 µm, 1.7 µm particle size, IonOpticks, Collingwood, VIC, Australia) coupled to a timsTOF Pro (Bruker, Billerica, MA) QTOF mass spectrometer.^38,41^ Mobile phase A was 0.1% formic acid in water and mobile phase B was 0.1% formic acid in ACN. The flow rate was 400 nL/min. The gradient consisted of a linear ramp from 2% B at 0 minutes to 17% B at 60 minutes, 25% B at 90 minutes, 37% B at 100 minutes, and 85% B at 110 minutes, which was held until 120 minutes.

Data were searched using DIA-Neural Network (DIA-NN, v1.9).^42^ A spectral library was generated using a human FASTA from Uniprot (canonical, accessed 03 May 2025). Trypsin/P was selected as the protease. The maximum number of missed cleavages was set to one, and the maximum number of variable modifications was set to two. Carbamidomethylation of cysteine was set as a fixed modification, and methionine oxidation and N-terminal acetylation were set as variable modifications. The output was filtered at 0.1% FDR.

### Lipidomics

Lipid extracts were dried using vacuum centrifugation and resuspended in 100 μL of 1:5 chloroform:MeOH. Condition-specific pooled samples, used for iterative MS^2^ analyses, were created by pooling 45 μL from each sample. Quality control (QC) samples were created by pooling 45 μL from each condition-specific pooled sample. Using a 1290 2DLC system (Agilent, Santa Clara, CA), 6 µL of sample was injected on a ZORBAX Eclipse Plus C18 column (2.1×100 mm, 1.8 µm, Agilent, Santa Clara, CA) in line with a 6545XT QTOF mass spectrometer (Agilent). Analyses were performed in both positive ion mode and negative ion mode. Mobile phase A consisted of 5:3:2 water:ACN:IPA with 10 mM ammonium acetate and 0.1% (v/v) InfinityLab Deactivator Additive (5191-4506, Agilent, Santa Clara, CA). Mobile phase B consisted of 90:9:1 IPA:ACN:water with 10 mM ammonium acetate. The gradient consisted of 15% B held for 2 minutes, a linear increase to 55% B at 8 minutes, 70% B at 22 minutes, 93% B at 23 minutes, 97% B at 27 minutes, and 100% B at 28 minutes, which was held until 31 minutes before returning to initial conditions at 31.4 minutes to equilibrate the column until 37 minutes. The flow rate was 200 µL/min. Details about instrument parameters are included in Supplementary Methods.

Spectral library searching was performed in MS-DIAL^43^ (v5.5.250820). Iterations from separate conditions were searched separately. MZmine^44^ (v4.6.1) was used for feature extraction, retention time correction, and normalization to the intensities of internal standards. Features were searched against a local database consisting of matches exported from MS-DIAL. Features whose retention time and mass error were within 0.25 minutes and 5 ppm, respectively, of an identified lipid were annotated.

### Metabolomics

Metabolomics extracts were resuspended in 60 µL of 7:2:1 ACN:water:MeOH. Condition-specific pooled samples were generated by combining 30 µL from each sample, and a pooled QC sample was prepared by combining 30 µL from each condition-specific pool. Using a 1290 2DLC system (Agilent, Santa Clara, CA), 6 µL of sample was injected on a zwitterionic hydrophilic liquid interaction chromatography (HILIC) column (InfinityLab Poroshell 120 HILIC-Z column, 2.1×150 mm, 2.7 µm, Agilent, Santa Clara, CA) coupled to a 6545XT (Agilent, Santa Clara, CA) QTOF mass spectrometer. Samples were analyzed in both positive and negative ion modes. Mobile phase A consisted of 20 mM ammonium acetate at pH 9.3 in water with 0.1% (v/v) InfinityLab Deactivator Additive (5191-4506, Agilent, Santa Clara, CA). Mobile phase B was pure ACN. The gradient conditions were as follows: 90% B held until 2 minutes, linear decrease to 78% B at 16 minutes, 60% B at 24 minutes, 10% B at 30 minutes, and held until 36 minutes, 90% B at 38.2 minutes, which was held until 46 minutes. The flow rate was 200 µL/min except between 38.2 and 44 minutes, where it was 250 µL/min to accelerate column re-equilibration. Instrument parameters are outlined in Supplementary Methods. Spectral library searching and feature extraction were performed as described for the lipidomics analysis.

### Data Analysis and Statistics

Data processing was performed using R (v4.3.2), and detailed descriptions of data processing, including filtering, normalization, and imputation, have been provided in Supplementary Methods. Proteomics data were filtered and imputed at the peptide level using *tidyproteomics* before aggregating to the protein level.^45–47^ A Gene Ontology

Subcellular Compartments (GOSCL) analysis of the top 200 most abundant proteins from sEVs isolated from MDA cells was performed using STRING.^48^ Proteomics results from PPT and SECUF samples were compared to UC samples using limma tests, and p-values were adjusted using the Benjamini-Hochberg method. Statistical significance required p_adj_ < 0.05 and |Log_2_(Fold Change| > 0.75. GOSCL analyses were performed to identify compartments implicated by proteins enriched in the PPT and SECUF conditions compared to UC. A list of all detected proteins was used as the background.

Metabolomics and lipidomics data were filtered against process blank samples, requiring a feature to be detected at least 3-fold greater intensity in samples compared to the process blank in two-thirds of samples for sEVs isolated from MDA cells and four-fifths of samples of at least one condition for plasma-derived sEVs. Due to polyethylene glycol contamination in the PPT samples, metabolic features were passed through a custom filter to remove features consistent with polyethylene glycol. After imputation and median alignment, positive and negative ion mode data were merged using MSCombine for both lipidomics and metabolomics datasets.^49^

Data integration was performed in MetaboAnalyst 6.0^27^ by uploading protein (UniProt) identifiers and small molecule (HMDB IDs) identifiers into the Joint Pathway Analysis modules to determine significantly enriched pathways (hypergeometric test, FDR < 0.05). Visualization of pathway coverage was performed using PaintOmics 4.0^26^ to demonstrate coverage of the “Endocytosis” KEGG pathway.^50^

## Results and Discussion

In this study, we developed a multi-omics platform, combining LC-MS-based proteomics, lipidomics, and metabolomics to characterize the molecular composition of sEVs. We demonstrated this platform using sEVs isolated from MDA cells using UC and then evaluated its clinical applicability using plasma-derived sEVs isolated by SECUF, PPT, and UC. These methods were selected because they represent three widely used but mechanistically distinct strategies for sEV isolation. UC separates vesicles primarily based on size and density through high-speed sedimentation and is commonly regarded as a reference method for obtaining relatively high-purity vesicle preparations, albeit with lower yield.^17,18,51^ SECUF isolates vesicles based on hydrodynamic size, enabling effective removal of soluble plasma proteins but with incomplete separation from similarly sized lipoproteins.^17,18,51^ In contrast, polymer precipitation concentrates vesicles through volume-exclusion effects that reduce vesicle solubility, resulting in high particle recovery but increased co-isolation of plasma proteins, lipoproteins, and other macromolecular components.^17,51^ Although multiple commercial precipitation kits are available, they rely on the same underlying polymer-based mechanism.^17,51^ Therefore, the PPT approach used here serves as a representative model of precipitation-based isolation methods. Recently, new approaches have been developed to isolate sEVs, including microfluidic devices, but have yet to be widely adopted.^52,53^ Therefore, we have selected these commonly utilized sEV purification approaches, UC, SECUF, and PPT. Comparing these complementary strategies enables systematic evaluation of how isolation mechanism influences sEV yield, purity, and downstream multi-omics profiles. In doing so, we show the broad applicability of the method and directly compare differences in sEV populations isolated using these different approaches (**Figure 1**).

**Figure 1.**
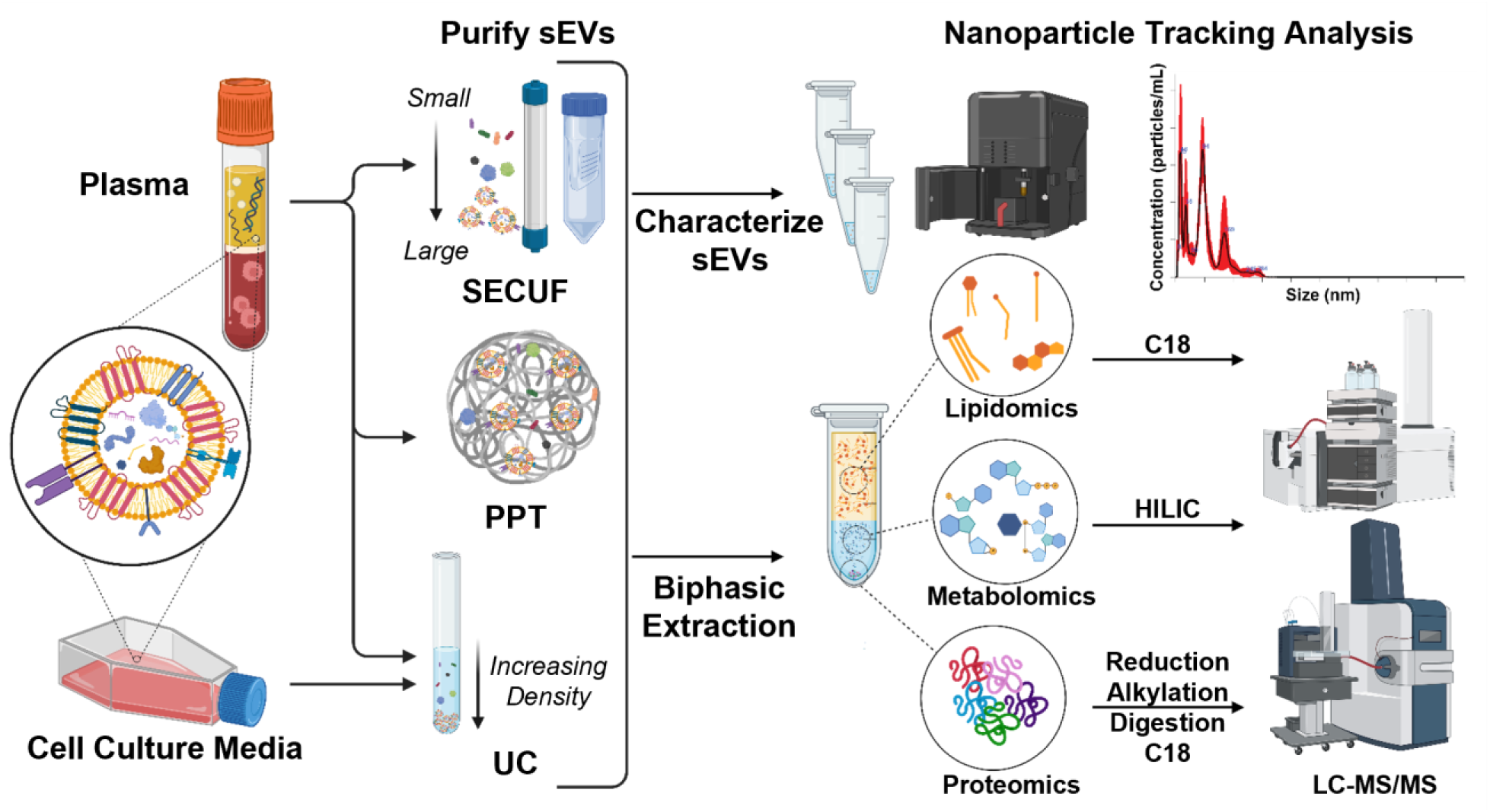
Schematic representation of the integrated multi-omics method for characterization of small extracellular vesicles (sEVs). Following nanoparticle tracking analysis (NTA), biphasic Matyash extraction enables sequential recovery of lipids, metabolites, and proteins from the same sEV sample. sEVs isolated from plasma by size exclusion chromatography with ultrafiltration (SECUF), polymer precipitation (PPT), and ultracentrifugation (UC) were analyzed to compare purification-dependent differences.

### Multi-omics of cell culture sEVs

To evaluate the performance of our multi-omics platform, 10 million sEVs, as determined by nanoparticle tracking analysis (**Figure S1**), were subjected to multi-omics analysis. Despite the limited sample input, we were able to achieve deep proteomics coverage, identifying an average of 43,152 peptides (**Figure 2A**) and 5,623 protein groups (**Figure 2B**, **Table S1**). The data were well normalized (**Figure S2A**) with low coefficients of variation (**Figure S2B**). Comparison of the identified proteins with databases of small extracellular proteins (ExoCarta^54^) and extracellular proteins (Vesiclepedia^55^) showed the majority of proteins detected (3,297) in this study are annotated in both databases (**Figure 2C**), with 67 proteins uniquely annotated in ExoCarta, 1,879 proteins uniquely annotated in Vesiclepedia, and 794 proteins annotated in neither. Furthermore, a GOSCL analysis of the top 200 most abundant proteins identified “Extracellular exosome”, “Extracellular membrane-bounded organelle”, and “Extracellular vesicle” as the top enriched terms (**Figure S2C**, **Table S2**), further indicating the detected proteins are relevant to sEVs.

**Figure 2.**
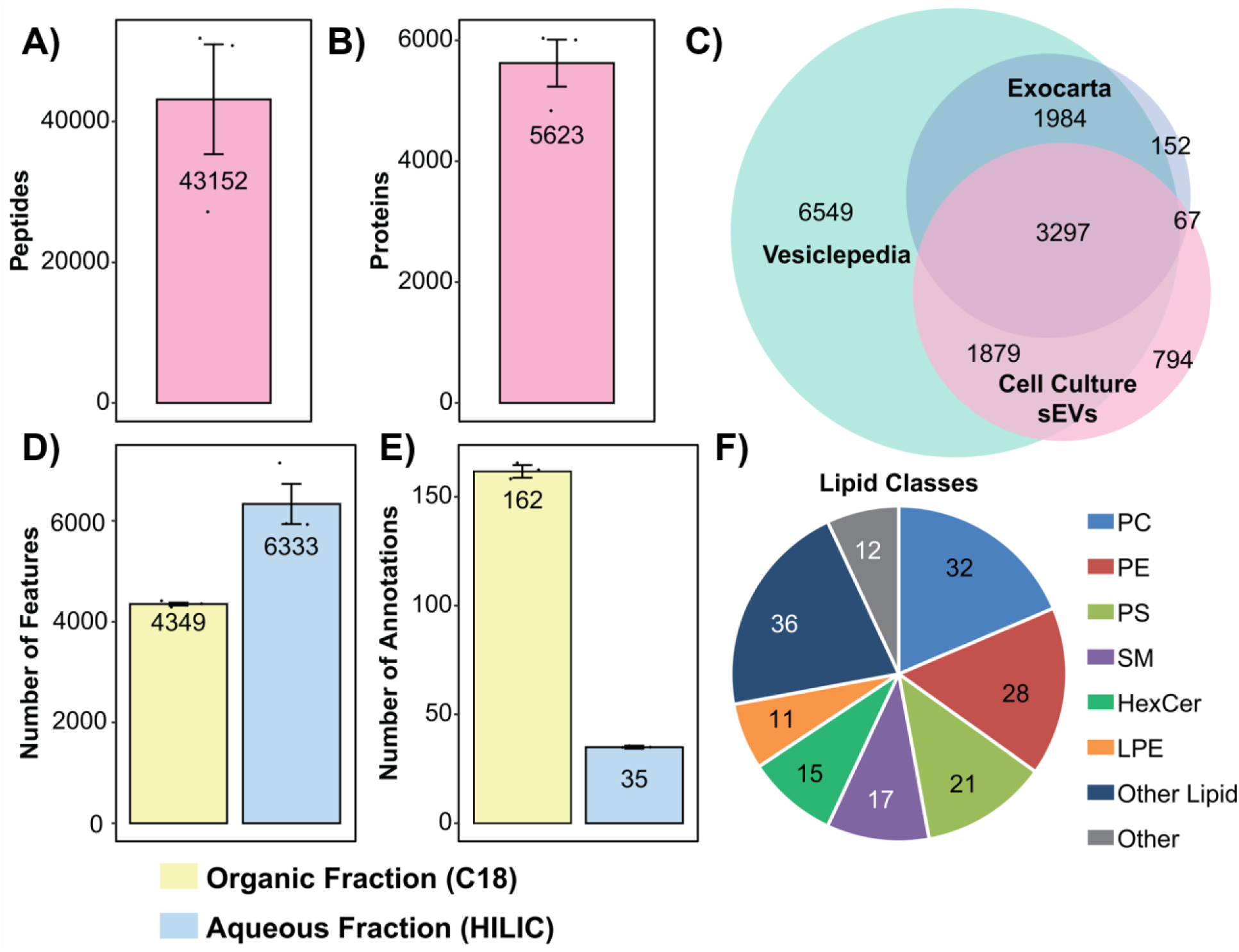
Multi-omics of sEVs isolated from MDA-MB-231 cells. Average number of (**A**) peptides and (**B**) protein groups identified from sEVs. (**C**) Overlap between proteins identified in sEVs isolated from conditioned cell culture media (pink) with databases of extracellular proteins (Vesiclepedia^55^; green) and small extracellular proteins (ExoCarta^54^; blue). (**D**) Average number of features found in the organic fraction (yellow) and aqueous fraction (blue). (**E**) Average number of annotated features in the organic fraction (yellow) and aqueous fraction (blue). (**F**) Lipid class distribution identified in the organic fraction. PC: phosphatidylcholines; PE: phosphatidylethanolamines; PS: phosphatidylserines; SM: sphingomyelins; HexCer: hexosylceramides; LPE: lysophosphatidylethanolamines.

In the analysis of the organic fraction (lipidomics), an average of 4,349 features were detected (**Figure 2D**), of which an average of 162 features were annotated (**Figure 2E**, **Table S3**). The consistent features and intensity of extracted ion chromatograms (EICs) of isotopically labeled internal standards (**Figure S3A-B)** and endogenous lipids (**Figure S3C-D**) from the lipidomics experiments indicate reproducible extractions and instrument performance in both positive and negative ion modes (**Figure S3**). The data are well normalized (**Figure S3E**) with a median coefficient of variation well below 5% (**Figure S3F**). When reviewing the lipidomic composition of the sEV population (**Figure 2F**), the majority of detected lipids were membrane-type phospholipids such as phosphatidylcholines (PC) and phosphatidylethanolamines (PE), similarly as reported previously.^56,57^ Only a small portion of detected lipids were triacylglycerols (TG) or cholesterol esters (CE), suggesting minimal lipoprotein contamination.^56–58^ In the analysis of metabolites extracted in the aqueous fraction (metabolomics), an average of 6,333 features were detected (**Figure 2D**), of which an average of 35 were annotated (**Figure 2E**, **Table S4**). Both the positive and negative ion mode EICs of internal standards and endogenous metabolites display reproducibility in intensity and retention time (**Figure S4A-D**), indicating reproducible extractions and proper instrument performance. This is further supported by consistency in the normalization (**Figure S4E**) and low coefficients of variation (**Figure S4F**). The relatively lower number of annotated polar metabolites (contained in the lumen) compared to lipids (contained in the membrane) likely reflects the high surface-to-volume ratio of sEVs.^59^ Previous studies have indicated that sEVs exhibit a propensity for containing their effector cargo within the vesicle membrane, in contrast to large EVs, which contain the majority of their cargo in the vesicle lumen.^59,60^

Integration of the multi-omics data was performed using Joint Pathway Analysis.^27^ The observed enrichment of pathways relating to breast cancer metabolism and proliferation aligns with the known tendency of sEVs to reflect the molecular state of their parent cells (**Figure 3**, **Table S5**).^4,6,11^ However, Joint Pathway Analysis identified “Endocytosis” as the most significantly enriched pathway (FDR < 1*10^-33^). Beyond sEV uptake, the endocytic pathway is of particular interest because exosomes are formed through an endosomal process in which intraluminal vesicles (ILVs) are created within multivesicular bodies.^3–7^ These multivesicular bodies dock with the luminal side of the plasma membrane, and ILVs are exocytosed as exosomes.^3–7^ Achieved coverage of the KEGG pathway, “Endocytosis”, is detailed in **Figure S5**, and many terms relating to the formation and maturation of endosomes were identified.^26,50^

**Figure 3.**
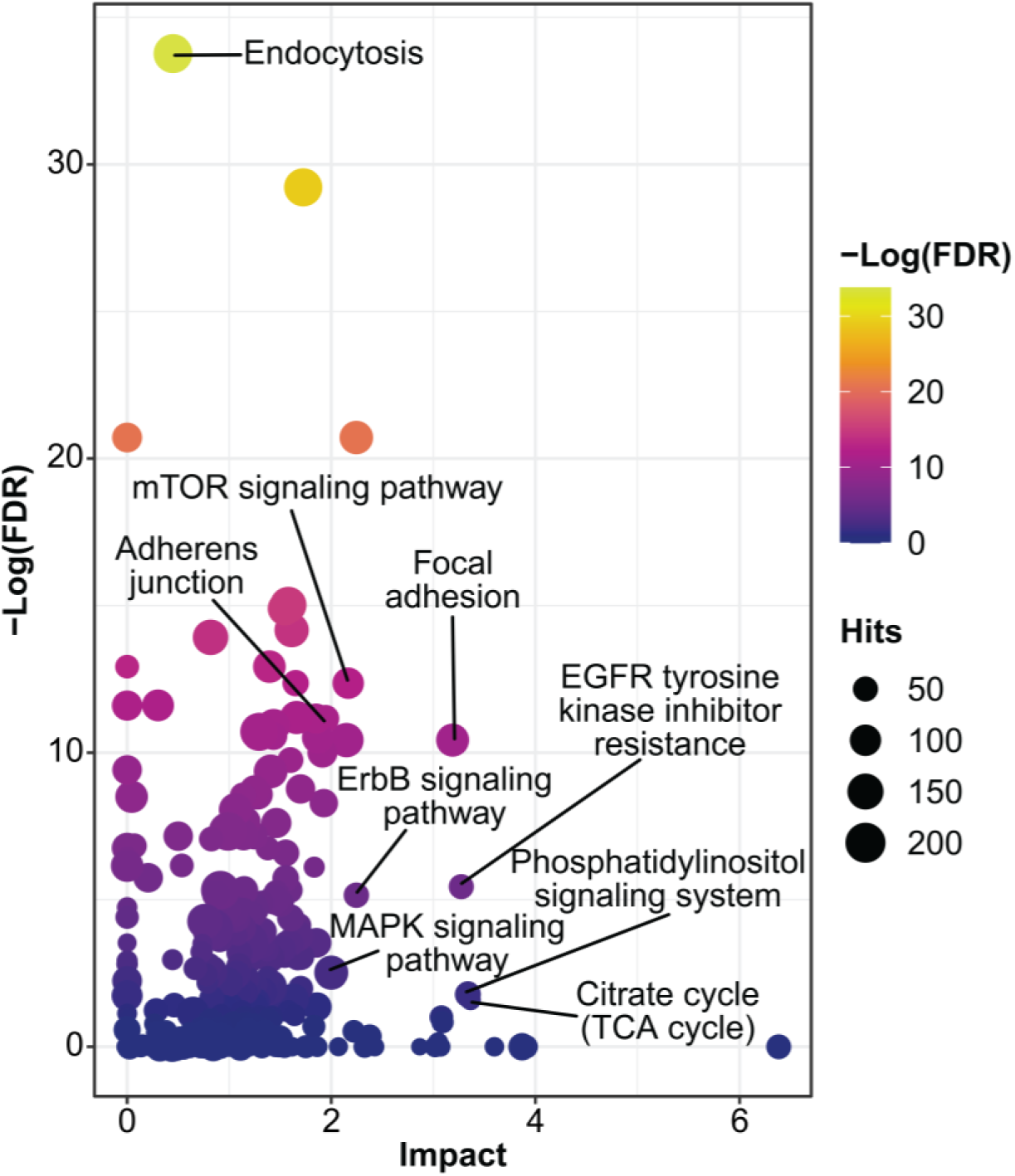
Joint pathway analysis of identified proteins and small molecules identified in cell culture-derived sEVs. Integrated multi-omics analysis were analyzed using the Joint Pathway Analysis module in MetaboAnalyst^27^ to identify significantly enriched pathways.

The results of this experiment indicate the ability of our method to achieve reproducible and relevant multi-omics characterization using as few as 10 million sEVs. Our analysis identified a broad range of proteins and lipids typical of sEVs, alongside key metabolites like purines (adenine, guanine, and hypoxanthine), and amino acids (methionine and arginine). Integration of the multi-omics data highlighted pathways linked to sEV biogenesis and release, as well as terms relevant to cancer biology, relating the molecular composition of sEVs to their parent cells.

### Purifying sEVs from Plasma

We next demonstrated the applicability of our method on sEVs isolated from plasma. To evaluate compatibility with our multi-omics platform, sEVs were purified from a single stock of human plasma using UC, SECUF, and PPT. Importantly, this experiment enables direct comparison of the molecular composition of the sEV populations isolated by different methods.

The data from nanoparticle tracking analysis of sEV populations resulting from UC, SECUF, and PPT purification from plasma are presented in **Figure S6**. For all three methods, nearly all particles larger than 200 nm were removed (**Figure S6A-C**), consistent with sEVs. Each method yielded a population of particles with an average diameter between 91 and 98 nm, and there is no significant difference in the average size of particle populations resulting from any of the purification approaches (**Figure S6D**). PPT had the greatest average yield of particles; greater than 10-fold more particles than SECUF and nearly 100-fold more particles than UC (**Figure S6E**). This trend is consistent with what has previously been reported in comparing these purification methods.^5,17^ Particle numbers were not standardized before extraction for multi-omics, as differences in yield are inherent to the purification approaches.^3,5,17,18^ Furthermore, the number of sEVs extracted influences the degree of small molecule coverage, and we aimed to provide an accurate depiction of sEV multi-omics coverage that is achievable when starting with 200 μL of human plasma. To enable comparative analysis of proteomes resulting from different extraction approaches, peptide loading was standardized to 250 ng. However, lipid and metabolite loading were not standardized; therefore, comparisons across purification methods focus on feature coverage and composition rather than relative intensity.

### Plasma sEV Proteomes Reflect Purification Approach

The proteomics analysis identified an average of 7,156 peptides from UC purified sEVs, 10,787 peptides from SECUF purified sEVs, and 5,954 peptides from PPT purified sEVs (**Figure 4A**). Correspondingly, an average of 1,260 protein groups were identified in UC samples, 1,582 proteins in SECUF samples, and 959 proteins in PPT samples (**Figure 4B**, **Table S6**). The reduction in proteomics coverage relative to sEVs isolated from MDA cells is attributable, at least in part, to abundant plasma proteins that co-purified with sEVs and obscured the detection of less abundant proteins.^5,17,18,61^ SECUF has improved performance in separating sEVs from blood proteins compared to UC and PPT, and this contributed to the increased proteomic coverage.^17,18,61^ Blood protein contamination was most pronounced in PPT-purified sEVs, which, despite having the highest protein yield (determined by Bradford assay following resolubilization of the protein pellet), exhibited the lowest proteomics coverage.

**Figure 4.**
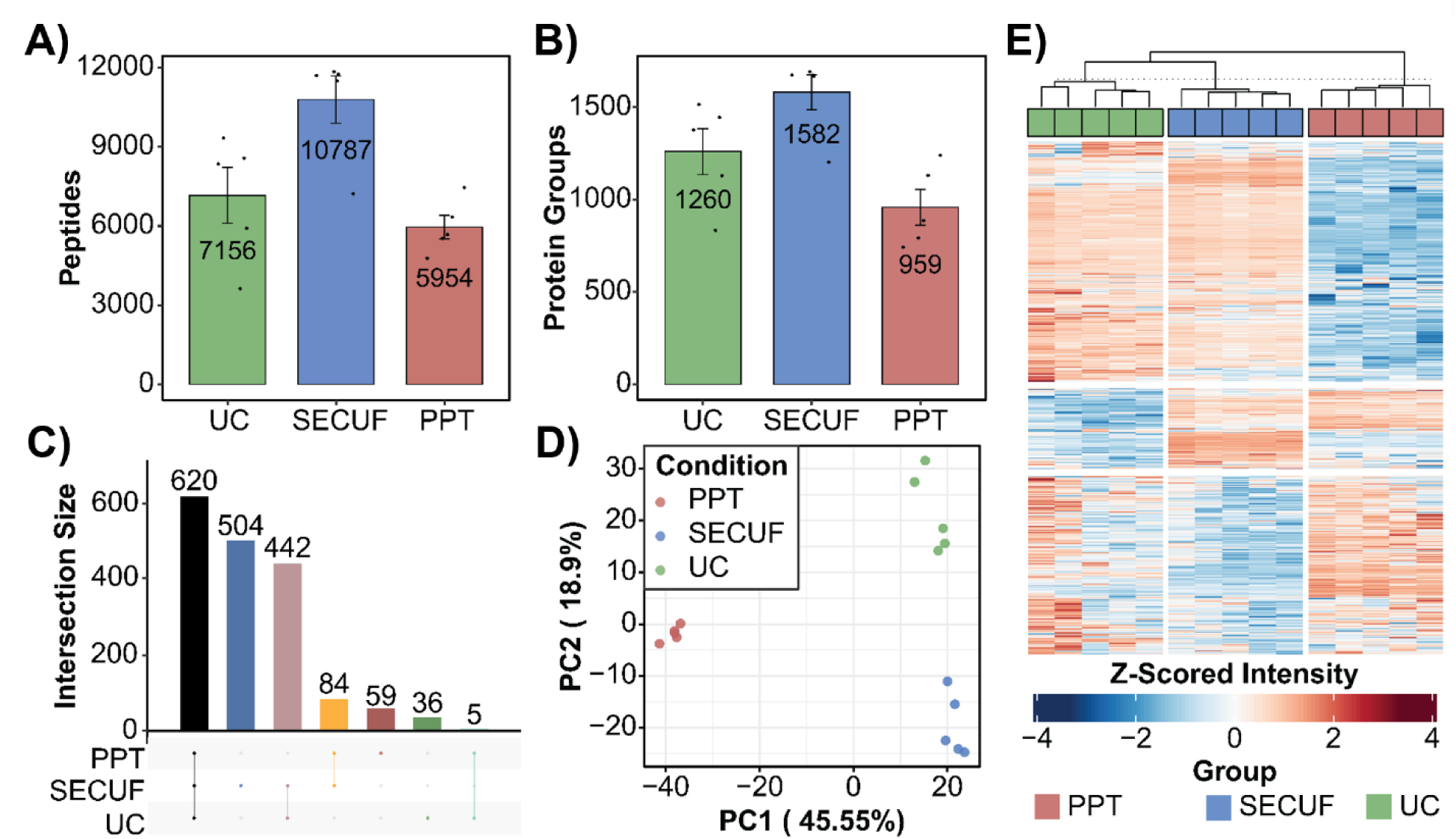
Plasma sEV proteomes reflect the purification approach. Average number of (**A**) peptides and (**B**) protein groups identified following ultracentrifugation (UC), size exclusion chromatography with ultrafiltration (SECUF), and polymer precipitation (PPT). (**C**) UpSet plot displaying the overlap in protein groups detected in each of the conditions. (**D**) Principal component analysis and (**E**) hierarchical clustering analysis illustrating the difference among UC, SECUF, and PPT.

Normalization among groups showed similar median intensities for UC and SECUF purified samples (**Figure S7A**). The median intensities of the PPT samples were lower which is likely due to the signal suppression from high-abundance proteins.^17,18,61^ Regardless of purification technique, the data were reproducible, with median coefficients of variation well below 5% within each group (**Figure S7B**). A total of 620 proteins were consistently detected across all purification methods, while more than 500 unique proteins were detected from the SECUF-purified sEVs (**Figure 4C**). Principal component analysis (PCA) indicates that sEV proteomes depend on the purification approach (**Figure 4D**). PPT purified proteomes are most distinct from SECUF and UC, separating out on the first principal component (PC1, 45.55%), whereas SECUF and UC separate on the second principal component (PC2, 18.9%). Hierarchical clustering analysis further supports this finding, where the PPT group is most unique, and SECUF and UC groups remain differentiated but are more similar (**Figure 4E**).

To assess the influence of the purification approach on the proteome, PPT and SECUF proteomes were compared against UC proteomes as UC is the standard purification method.^3,5,17–21,58,62^ PPT purification resulted in significantly lower (p_adj_ < 0.05, Log_2_(Fold Change) < -0.75) relative levels of TSG101, CD81, and CD9 (**Figure 5A**), which are sEV biomarkers.^3^ CD63, another sEV biomarker,^3^ was also lower in the PPT condition but failed to reach our Log_2_(Fold Change) cutoff for significance (p_adj_ = 0.010, Log_2_(Fold Change) = -0.67). The lower abundance of sEV biomarkers indicates lower purity sEV populations compared to UC. In addition, the intensity of APOB, a biomarker of low-density lipoproteins (LDL),^17,18^ is significantly increased in PPT samples. However, there was no significant difference in the abundance of LPA, a biomarker of high-density lipoproteins (HDL).^17^ In a GOSCL analysis of proteins enriched in PPT samples (using all detected proteins as background), three of five enriched terms are related to lipoproteins, providing additional evidence of lipoprotein contamination (**Figure 5B**, **Table S7**).

**Figure 5.**
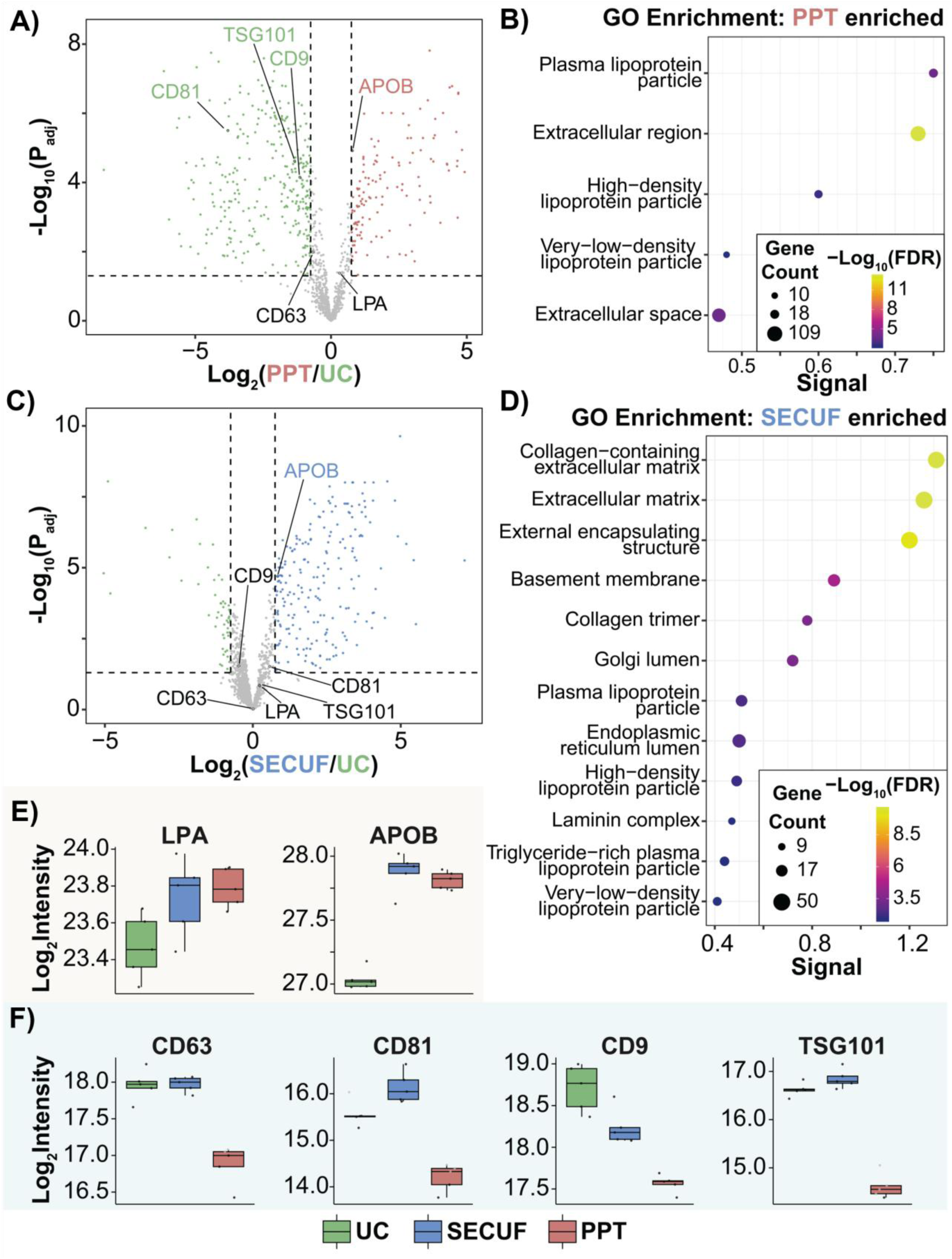
Comparative proteomics indicate differences in the purity of sEV preparation approaches. (**A**) Volcano plot illustrating proteomic differences between sEV populations isolated by ultracentrifugation (UC, green) and polymer precipitation (PPT, red). (**B**) Gene Ontology Subcellular Localization (GOSCL) analysis of proteins that are significantly more abundant in PPT compared to UC. (**C**) Volcano plot illustrating proteomic differences in sEV populations isolated using UC (green) and size exclusion chromatography with ultrafiltration (SECUF, blue). (**D**) GOSCL analysis of proteins that are significantly more abundant in SECUF proteomes compared to UC. (**E**) Box-and-whisker plots showing the differences in the abundances of lipoprotein-associated proteins across purification conditions. (**F**) Box-and-whisker plots showing the differences in the abundances of canonical sEV biomarker proteins across purification approaches.

Comparison of UC and SECUF sEV proteomes showed no significant difference in CD63, CD81, CD9, or TSG101 abundances (**Figure 5C**), indicating comparable enrichment of vesicle-associated proteins. However, APOB was significantly more abundant in SECUF samples compared to UC, suggesting LDL contamination in the SECUF sEV population. This finding is consistent with previous literature indicating LDL contamination in sEVs purified using size exclusion chromatography (SEC).^5,18^ Furthermore, GOSCL analysis of proteins significantly more abundant in SECUF purified sEVs highlights terms related to the extracellular matrix and lipoproteins, among others (**Figure 5D**, **Table S8**).

Overall, UC purified sEVs had the lowest abundance of LPA and APOB, although LPA did not differ significantly between UC and either PPT or SECUF (**Figure 5E**). PPT purified sEVs had the lowest abundance of sEV biomarkers, while comparable intensities were observed in the UC and SECUF sEV populations (**Figure 5F**). These data suggest that the PPT-purified sEV population suffers from the greatest contamination. The similar relative abundance of sEV biomarkers between UC and SECUF purified sEVs indicates similar overall purities, but the higher abundance of APOB suggests more LDL contamination in the SECUF condition.

### Purification Approach Affects Purity and Composition of the sEV Metabolome

Lipidomics coverage varied depending on the purification approach. The fewest number of features was detected in the UC group, followed by SECUF, and the greatest number were observed in the PPT condition (**Figure 6A**, **Table S9**). A similar trend was observed for the annotated features. The fewest number of features were annotated in the UC condition (an average of 142), with an average of 440 annotated features in the SECUF condition, and 837 annotated features in the PPT condition (**Figure 6B**). These variations are driven primarily by differences in material yield across the purification methods, and secondarily by differences in composition of the resultant population.

**Figure 6.**
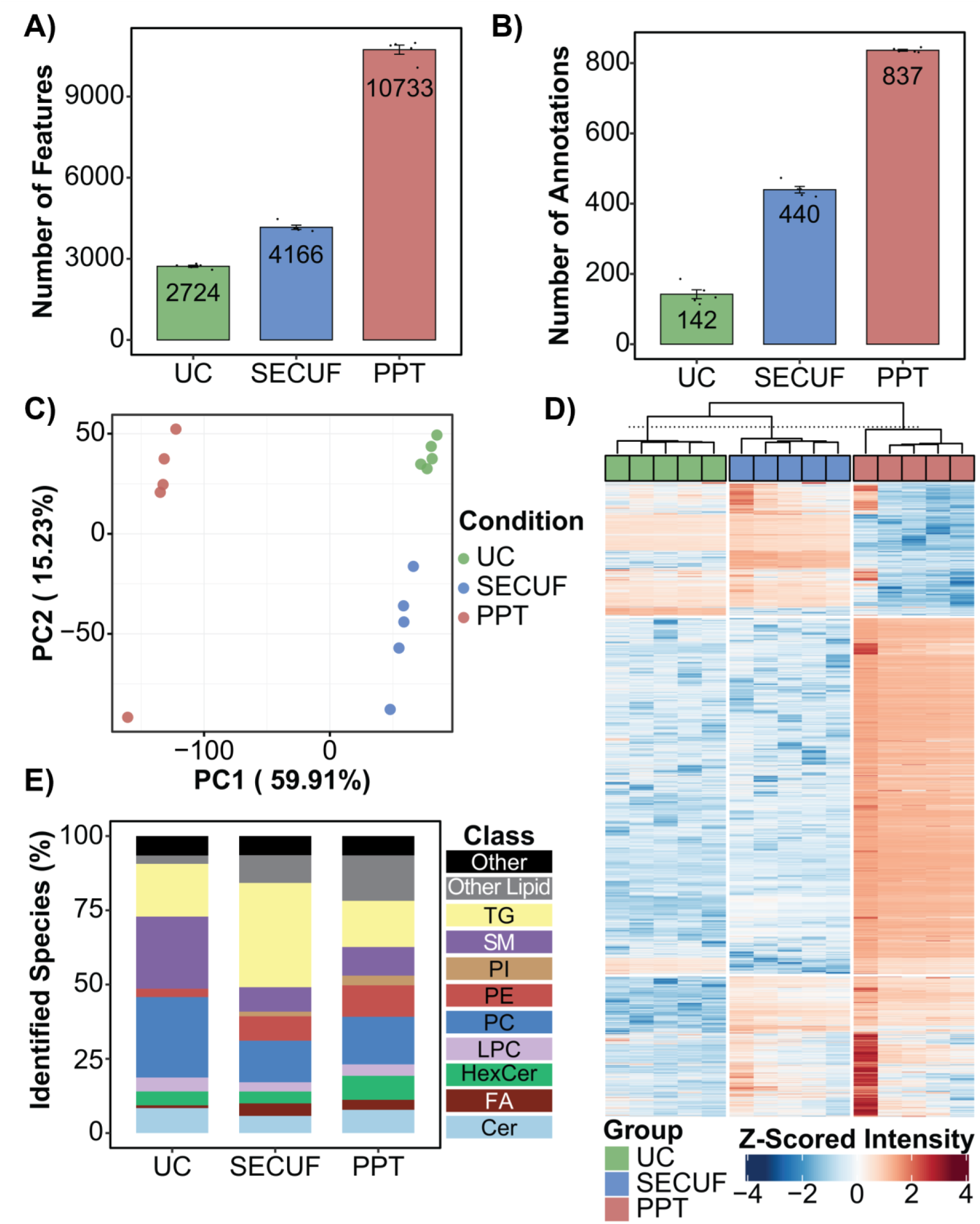
Lipidomics analysis of sEVs from plasma reveals population differences associated with purification approach. Average number of (**A**) features and (**B**) annotated features identified through analysis of the organic fraction of extracted sEVs purified using ultracentrifugation (UC), size exclusion chromatography with ultrafiltration (SECUF), and polymer precipitation (PPT). (**C**) Principal component analysis and (**D**) hierarchical clustering illustrating the dissimilarity among the different purification approaches. (**E**) Stacked bar plot illustrating the relative lipid class composition of annotated features in the sEV lipidomes across purification approaches. Cer: ceramides; FA: fatty acids; HexCer: hexosylceramides; LPC: lysophosphatidylcholines; PC: phosphatidylcholines; PE: phosphatidylethanolamines; PI: phosphatidylinositols; SM: sphingomyelins; TG: triacylglycerols.

The EICs of isotopically labeled internal standards showed stable retention times and consistent intensities within each condition (**Figure S8A-B**). However, there is variability between samples purified with different techniques, which is attributable to ion suppression from different amounts of co-eluting species and matrix effects on extraction efficiency arising from differences in the amount and composition of material extracted.^63^ Data were normalized in each group (**Figure S8C-E**), but the median feature intensities varied depending on purification technique, as it was highest for samples purified using PPT, and lowest for samples purified with UC. This difference is attributable to differences in the amount of material injected on the column. High precision was achieved in each group, with the median coefficient of variation less than 5% in each condition (**Figure S8F-H**). As was seen in the proteomics analysis, the PPT group was most distinct from the UC and SECUF group, separating out on PC1 (59.91%) in a PCA (**Figure 6C**). The UC and SECUF conditions were also unique, separating on PC2 (15.23%). The hierarchical clustering analysis further supports this trend, with the PPT condition separating from SECUF and UC, which remain distinguishable from each other (**Figure 6D**).

Analysis of the relative lipidomic composition (**Figure 6E**) revealed that the UC condition had the highest relative composition of membrane-type phospholipids, including sphingomyelins (SM), PEs, PCs, and lysophosphatidylcholines (LPC), consistent with vesicle membrane composition. TGs constituted a significant proportion of identified lipids regardless of purification approach, but they accounted for the highest proportion of lipids identified in SECUF purified sEV populations, as a result of LDL contamination.^5,17,18^ Although similar lipid classes account for the majority of annotated features across purification methods, only 59 lipids were identified in all three conditions (**Figure S8I**), with an additional 280 lipids shared between PPT and SECUF conditions.

Metabolomic results showed trends similar to lipidomics. The fewest number of features were detected in the UC condition (3,366), followed by SECUF (4,022), with the greatest number of features identified in the PPT condition (20,173) (**Figure 7A**, **Table S10**). Accordingly, the fewest average number of features were annotated in the UC sEV populations (34), with the SECUF condition having the next highest average (48), and the greatest number of average annotated features were found in the PPT condition (208) (**Figure 7B**). EICs of internal standards (**Figure S9A-B**) display consistency in retention time and intensity across most samples, but some SECUF samples exhibit inconsistent retention time and abundances of standards, likely due to the salt carryover from SEC buffers. The high concentration of salt contributed to ion suppression and interacted with the zwitterionic HILIC column, affecting analyte retention.^64^ Nevertheless, within each group, the data had similar median normalized abundance values (**Figure S9C-E**). In addition, the median coefficient of variation was below 5% in each condition (**Figure S9F-H**). PCA of metabolomics data (**Figure 7C**) showed similar results to what was observed in the lipidomics and proteomics datasets. The PPT condition separates from the UC and SECUF conditions on PC1 (57.06%), and the UC and SECUF conditions separate on PC2 (18%). These results are further supported by the hierarchical clustering analysis (**Figure 7D**), where the PPT samples are most unique, and the SECUF and UC samples are more similar but can still be separated from each other. The majority of annotated metabolites were unique to the condition in which they were detected (**Figure 7E**).

**Figure 7.**
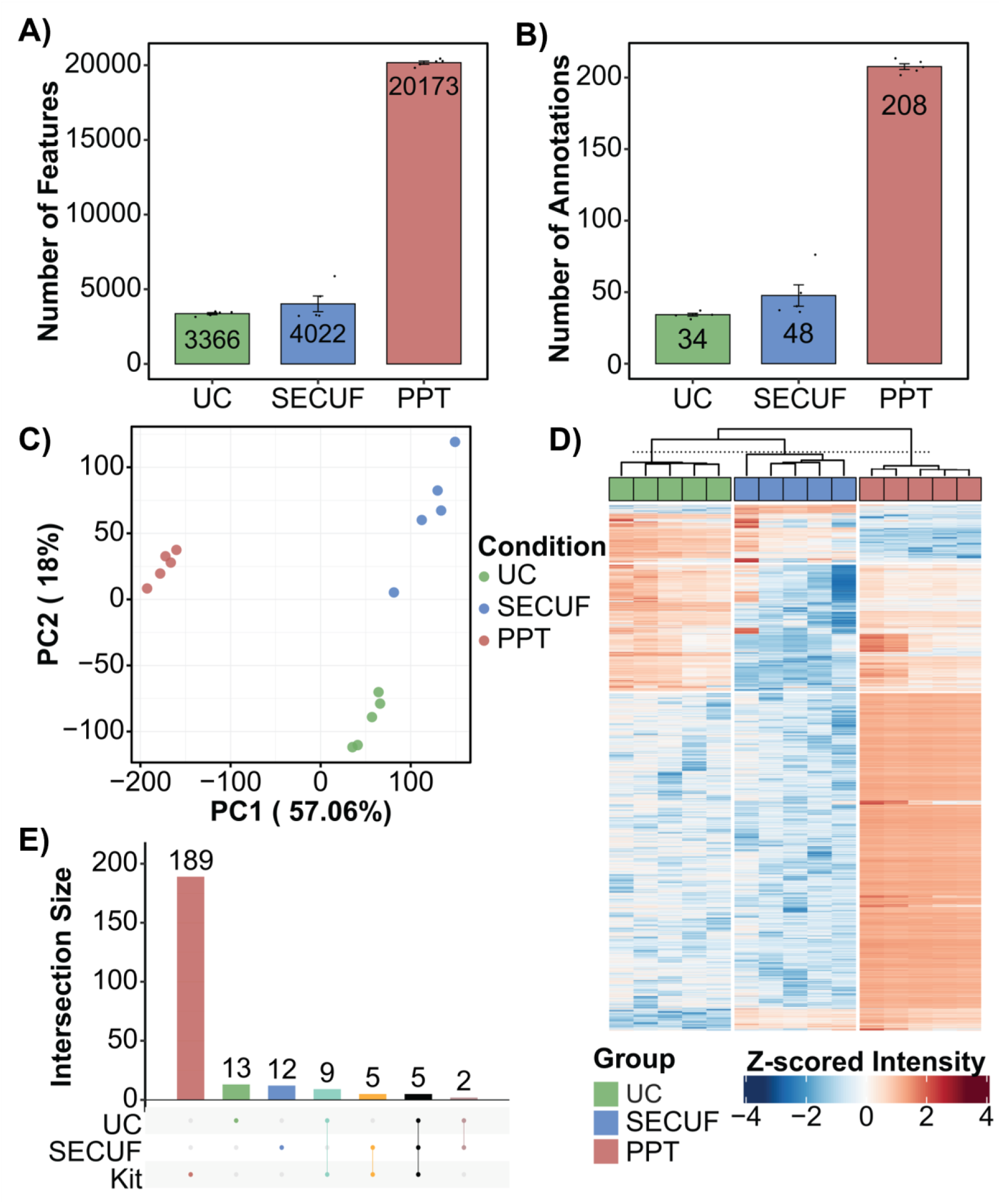
Metabolomic coverage and composition depend on the purification approach. Average number of (**A**) features and (**B**) annotated features identified using ultracentrifugation (UC), size exclusion chromatography with ultrafiltration (SECUF), and polymer precipitation (PPT). (**C**) Principal component analysis illustrating differences in the metabolome among the different purification approaches. (**D**) Hierarchical clustering analysis showing the difference across purification methods. (**E**) UpSet plot displaying the degree of overlap in identified features among the different conditions.

While our multi-omics platform was successfully applied across all three isolation methods, sEVs purified via SECUF and PPT introduced inherent technical challenges that complicated metabolomics analysis. In SECUF purified sEVs, the salts required for SEC separations had a deleterious effect on both analyte retention and detection. For PPT purified sEVs, polymer contamination resulted in ion suppression and additional filtering during data analysis.^65^ UC purified sEVs were most compatible with our method, and deeper metabolomic coverage may be achieved with increased plasma input.

### Integrated Analysis Reveals Purification Method-Dependent Multi-omics Signatures of sEVs

The multi-omics data were integrated using Joint Pathway Analysis of identified proteins and small molecules (**Tables S11-S13**). Across all purification methods, “Endocytosis” was a significantly enriched term (**Figure S10A-C**), consistent with the endosomal origin and release of exosomes. In addition, pathways related to interactions with the extracellular space were significantly enriched for each approach, including “ECM-receptor interaction” and “Focal adhesion”. The most significantly enriched pathway, regardless of purification method, was “Complement and coagulation cascades”, likely reflecting the co-purification of high-abundance plasma proteins with the sEVs.^17,18,61^ This term showed the highest enrichment in the PPT condition, while “Endocytosis” showed the lowest. This further suggests that sEVs purified via PPT maintain the lowest level of purity. Furthermore, the fewest number of significantly enriched pathways were identified in the PPT condition (45), whereas 80 and 97 were identified in the UC and SECUF conditions, respectively. Out of the 100 unique significantly enriched pathways, 81 were common to at least two conditions, of which 41 were enriched across all three methods (**Figure S10D**).

Our results indicate the tradeoffs in purity and yield across sEV purification methods. UC yields the highest purity sEV population,^18^ but is limited by low recovery and a time-consuming purification process. Despite maintaining the highest purity, it is impossible to remove all contaminants with UC, as HDL and plasma proteins can co-precipitate with sEVs.^17,18^ SECUF is more effective in separating plasma proteins from sEVs,^17,18^ and provides a higher particle yield, resulting in greater proteomics coverage. Moreover, in our analysis, the relative abundances of several sEV biomarkers were comparable to what was observed using UC, indicating similar purity; however, the salt introduced for SEC separations makes this approach less compatible with metabolomics analysis. Furthermore, as LDL particles and sEVs are similar in size, LDL contamination is a significant challenge in SEC-purified sEVs.^5,18^ This contamination was evident in our analysis by increased abundances of APOB, and a greater proportional representation of TG. PPT yielded the largest amount of particles (based on NTA) and proteins (based on the protein assay), but the low abundance of sEV biomarkers and greater abundance of lipoprotein-related proteins indicate substantial non-sEV contamination. Furthermore, the residual polymer drastically complicates metabolomics analyses. Taken together, our data indicate that UC is still the optimal method for purifying sEVs for multi-omics analysis, balancing high sEV purity with minimal introduction of interfering species. We anticipate that the starting volume of plasma, and subsequently, the sEV yield, will be a key determinant of achievable molecular coverage.

## Conclusion

We developed an integrated multi-omics platform that enables the characterization of as few as 10 million sEVs. Our strategy maximizes sample utilization through sequential extraction of lipids, metabolites, and proteins from the same sample. Metabolomics and lipidomics experiments were tailored to maximize quantitative precision and molecular coverage by implementing an iterative MS^2^ strategy. Deep proteomics coverage was achieved by performing nanoflow separations with diaPASEF. We demonstrated this method on sEVs purified from MDA cells, and identified an average of 5,623 proteins and 197 lipids or metabolites, providing a holistic, integrated molecular characterization. The quantitative robustness and molecular coverage of this method support its potential utility for the identification of biomarkers, and we further demonstrated its compatibility with sEVs isolated from human plasma using UC, SECUF, and PPT. We identified molecular differences in sEV populations, reflecting inherent tradeoffs in sEV purity and yield that affect the multi-omic profile: UC produced the highest vesicle purity, PPT yielded the greatest particle recovery but the lowest purity with increased plasma and lipoprotein contamination, and SECUF showed intermediate characteristics with preserved vesicle markers but detectable LDL carryover. We envision this integrated multi-omics platform will enable future biomarker and functional studies by providing an integrated view of the protein, lipid, and metabolite cargoes in single sEV samples.

## Acknowledgments

This work is supported by the University of Wisconsin - Madison Office of the Vice Chancellor for Research with funding from the Wisconsin Alumni Research Foundation (to Y. G., S. J., and S. M. P.). We would like to thank Dr. Alissa Weaver for providing the MDA-MB-231 cell line stably expressing mScarlet-CD63. H.T.R. would like to acknowledge support from the National Heart, Lung, and Blood Institute of the NIH under Award Number T32HL007936 through the UW-Madison Cardiovascular Research Center and NIH F31HL178305.

