## Supporting Information for "Integrated Multi-Omics Enabled by Sequential Extraction for Comprehensive Molecular Profiling of Small Extracellular Vesicles"

### Table of Contents


**Supplementary Methods**

Purification of Small Extracellular Vesicles (sEVs) from MDA-MB-231 cells using serial ultracentrifugation……..………………........................................S-03

Purification of sEVs from plasma………………… ……………………………S-04

Instrument parameters………………………… …………………………… S-06

Data Analysis and Statistics………………………….…………………………..S-08

**Supplementary Figures**

**Figure S1.** Nanoparticle tracking analysis of small extracellular vesicles (sEVs) purified from conditioned cell culture media using ultracentrifugation (UC)..………………………………………………………………………….S-10

**Figure S2.** Proteomics analysis of small extracellular vesicles (sEVs) isolated from conditioned cell culture media……………………………………...……… S-11

**Figure S3.** Lipidomics analysis of small extracellular vesicles purified from conditioned cell culture media……...…………………………….……………… S-12

**Figure S4.** Metabolomics analysis of small extracellular vesicles (sEVs) isolated from conditioned cell culture media……………..………..……………………. S-14

**Figure S5.** Coverage of molecules implicated in the endocytosis pathway.. S-16

**Figure S6.** Nanoparticle tracking analysis of small extracellular vesicles (sEVs) isolated from plasma using ultracentrifugation (UC), size exclusion chromatography with ultrafiltration (SECUF), and polymer precipitation (PPT).……………………..………………………………………………….. S-17

**Figure S7.** Proteomics of small extracellular vesicles (sEVs) purified from plasma using ultracentrifugation (UC), size exclusion chromatography with ultrafiltration (SECUF), and polymer precipitation (PPT)…………………………….……….. S-19

**Figure S8.** Lipidomics analysis of small extracellular vesicles purified from plasma using ultracentrifugation (UC), size exclusion chromatography with ultrafiltration (SECUF), and polymer precipitation (PPT)……………………………..……..… S-20

**Figure S9.** Metabolomics analysis of small extracellular vesicles isolated from plasma using ultracentrifugation (UC), size exclusion chromatography with ultrafiltration (SECUF), and polymer precipitation (PPT)……………………S-22

**Figure S10.** Joint pathway analysis of circulating small extracellular vesicles……………………………………………………………………………S-24

**References**……………………………………………………………………….S-25

**Supplementary Methods.**

**Purification of Small Extracellular Vesicles (sEVs) from MDA-MB-231 cells using serial ultracentrifugation.**

MDA-MB-231 cells stably expressing mScarlet-CD63 (a kind gift from Dr. Alissa Weaver) were cultured in DMEM supplemented with 10% FBS.[1] Upon reaching 70-80% confluency, the standard medium was replaced with Opti-MEM to minimize contamination from bovine serum EVs, and the cells were incubated for 48 hours. The conditioned media were collected for sEV isolation using differential ultracentrifugation. First, the media was centrifuged at 300 × *g* for 10 minutes to remove cells, followed by a second spin at 2,000 × *g* for 20 minutes to eliminate dead cells and debris. Finally, to pellet and remove microvesicles, the collected supernatant was centrifuged at 10,000 × *g* for 30 minutes at 4°C, and the remaining supernatant containing exosomes was transferred to 50 mL fresh tubes. Then, the cell media containing CD63-mScarlet-labeled sEVs was concentrated to a final volume of 12 mL using a Centricon® Plus-70 device (Sigma-Aldrich). The concentrated media was transferred into ultracentrifuge tubes (Ultra-centrifuge tube 344060, Beckman Coulter) and centrifuged at 100,000 × *g* for 4 hours at 4 °C. The pellets were resuspended in DPBS and spun again in the same conditions. The final pellet containing purified sEVs was resuspended in DPBS for characterization of sEV number and size measured by NanoSight-NS300 nanoparticle tracking analysis (NTA) (Malvern Panalytical) and Western blot analysis. Based on the NTA results, 10 million MDA-MB-231-derived sEVs were aliquoted for multi-omics analysis.

#### Purification of sEVs from Human Plasma

Upon receipt, plasma was stored at -80 °C. Once thawed, plasma was pooled to create a sample with sufficient volume for all experiments. The pooled plasma was centrifuged at 2,000 × *g* at 4 °C for 20 minutes to remove cell debris. The resulting supernatant was aliquoted out into 1.5 mL tubes (200 μL/sample) and stored at -80 °C until sEV purification.

Polymer precipitation was performed using the Total Exosome Isolation (from plasma) kit (Invitrogen, Waltham, MA) without Proteinase K treatment. The decellularized plasma was thawed on ice and then centrifuged at 18,000 × *g* at 4 °C for 30 minutes to remove large vesicles. Supernatants were transferred to new 2.0 mL tubes, and 100 μL of Dulbecco's Phosphate Buffered Saline (DPBS) was added to each sample and mixed by vortexing. Sixty μL of the precipitation reagent was added to each of the samples, which were vortexed briefly and incubated at room temperature for 10 minutes. Subsequently, the samples were centrifuged at 10,000 × *g* for 10 minutes at room temperature, and the supernatant was aspirated away. Centrifugation and supernatant removal were repeated. The purified sEVs were resuspended in 100 μL of DPBS and snap frozen for characterization using nanoparticle tracking analysis (NTA) and multi-omics analysis.

For SECUF purification, decellularized plasma was thawed on ice, and large EVs were removed via centrifugation at 18,000 × *g* at 4 °C for 30 minutes. The supernatants were transferred to new 2.0 mL tubes and passed through a qEVoriginal 35 nm Gen 2 SEC column (Izon, Christchurch, New Zealand). To equilibrate the column, it was flushed with 17 mL of DPBS. The 200 μL sample volume was loaded on the frit, and buffer volume collection began immediately. Once the sample entered the column, DPBS was loaded on the frit as the mobile phase. After the buffer volume was collected (2.5 mL), the next 1.2 mL of elution volume was collected. An additional 1.6 mL of DPBS was run through the column before rinsing with 8.5 mL of 0.5 M NaOH and flushing with 17 mL of DPBS. After flushing, the next sample was loaded. The samples were concentrated to less than 200 µL using 30 kDa molecular weight cut-off filters (MWCO) (REF #UFC903008, Millipore, Burlington, MA). The concentrate was transferred to clean 2.0 mL tubes, snap frozen, and stored at -80 °C for NTA and multi-omics analysis.

For ultracentrifugation, decellularized plasma was thawed on ice and centrifuged at 18,000 × *g* for 30 minutes at 4 °C to remove microvesicles. Supernatants were transferred to ultracentrifuge tubes (344057, Beckman Coulter, Brea, CA) and diluted to 3 mL with DPBS. Samples were centrifuged at 100,000 × *g* at 4 °C for 2 hours. The supernatants were removed, and the pellets were resuspended with 750 µL of cold DPBS, transferred to a new ultracentrifuge tube, and diluted to 3 mL with DPBS. The centrifugation step was repeated, the supernatants were removed, and the pellets were resuspended in 100 µL of cold DPBS before being transferred to 2.0 mL tubes. The resuspended sEVs were snap frozen and stored at -80 °C for NTA and multi-omics analysis.

**Instrument parameters.**

**Proteomics.** Proteomics analyses were performed on a timsTOF Pro operated in the positive ion mode (PIM) using data independent analysis parallel accumulation serial fragmentation (diaPASEF).[2] The MS scan range was 100-1700 m/z. The capillary voltage was 1650 V, and the dry gas temperature and flow rate were 181 °C and 3 L/min, respectively. diaPASEF was performed using 32 windows ranging from m/z 400 to 1201 and $1/K_0$ 0.6 to 1.43.

**Lipidomics.** For lipidomics, liquid chromatography-mass spectrometry (LC-MS) analysis was performed using an Agilent 1290 2DLC in line with a 6545XT Q-TOF mass spectrometer. The second-dimension pump was used to deliver 50 µL/min of TOF reference mass, consisting of 5 µM purine and 1.25 µM Hexakis(1H,1H,3H-tetrafluoropropoxy)phosphazine in 95% ACN. To maximize sensitivity and coverage, samples were analyzed using $MS^1$ only, and condition-specific pooled samples were analyzed using an iterative $MS^2$ approach consisting of 5 iterations in each ion mode for spectral library search-based annotations.[3] The gas temperature, drying gas flow, nebulizer pressure, sheath gas temperature, and sheath gas flow were 200 °C, 10L/min, 35 psi, 300 °C, and 12 L/min, respectively. The capillary voltage was 3500 V and the Nozzle Voltage was 1000 V in PIM, and they were 3000 V and 1500 V, respectively, in the negative ion mode (NIM). Fragmentor, Skimmer, and Octupole RF were set to 150 V, 65 V, and 750 Vpp, respectively. The mass range was 50-1700 m/z, and the instrument operated at 3 Hz in PIM and 1.5 Hz in NIM. $MS^2$ analyses were performed across the same mass range and $MS^2$ spectra were acquired at 2.25 Hz in PIM and 1.13 Hz in NIM, and collision energy was set to 20 V to induce collisionally activated dissociation (CAD).

**Metabolomics.** For metabolomics, LC-MS analysis was performed using a 1290 2DLC system (Agilent) in line with a 6545XT Q-TOF mass spectrometer. Reference mass was delivered using the second-dimension pump at a constant rate of 50 µL/min. Samples were analyzed using $MS^1$ only, and condition-specific pools were analyzed using an iterative $MS^2$ approach, consisting of 5 iterations in positive ion mode and 3 iterations in negative ion mode. The gas temperature, drying gas flow, nebulizer pressure, sheath gas temperature, and sheath gas flow were 225 °C, 9 L/min, 40 psi, 375 °C, and 12 L/min, respectively. The fragmentor, skimmer, and octupole voltages were 100 V, 45 V, and 750 Vpp, respectively. In both ion modes, the capillary voltage was 3000 V and the nozzle voltage was 500 V. The mass range was 50-1400 for $MS^1$, and 40-1100 m/z for $MS^2$. $MS^1$ scans were performed at 3 Hz in PIM and 1.5 Hz in NIM. For $MS^2$ analyses, 20 V of collision energy was applied, and $MS^2$ scans were collected at 2.25 Hz in PIM and 1.13 Hz in NIM.

**Data Analysis and Statistics**

Data processing was performed using R (v4.3.2). Proteomics data were filtered and imputed at the peptide level using *tidyproteomics* with modifications.[4–6] For the cell culture sEV experiments, peptides were retained if they were detected in at least two-thirds of the samples. For the plasma sEV experiments, peptides were retained if they were detected in at least four-fifths of samples in at least one condition. Intensity values were normalized using the random forest normalization implemented in *tidyproteomics*. Missing values were imputed using MissForest[7] unless values were missing entirely within a condition (MEC), in which case, they were imputed using quantile regression imputation of left-censored values (QRILC).[5] Peptide data were aggregated to the protein level using the collapse function in *tidyproteomics*. For comparative analyses, limma tests were performed, and p-values were adjusted using the Benjamini-Hochberg correction. Statistical significance required $p_{adj} < 0.05$ and $|Log_2(\text{Fold Change})| > 0.75$. Gene ontology analyses were performed using STRING to determine whether specific subcellular compartments are implicated by the proteins significantly enriched in PPT or SECUF compared to UC, using a list of all detected proteins as the background.

For the cell culture sEV experiments, metabolite and lipid features were filtered such that any feature that was missing or detected at less than three-fold the intensity of the process blank in more than one sample was removed. For the plasma sEV experiments, features were retained if they were detected at more than 3-fold the intensity of the process blank in at least four-fifths of samples in at least one condition. The plasma sEV metabolic features were subjected to a custom filter to remove features consistent with polyethylene glycol, introduced in the PPT purification. This filtering was performed by identifying and removing co-eluting features with mass differences consistent with the monomeric mass of polyethylene glycol. Imputation was performed using QRILC[5] for MEC values and MissForest[7] for all other values. Positive and negative ion mode data frames were concatenated using MSCombine, following median alignment.[8] Common features were identified by considering common adducts, requiring a difference in retention time of less than 0.25 minutes, a difference in mass (having considered adducts) of less than 10 ppm, and correlated feature intensities within samples between ion modes.

#### Supplementary Figures

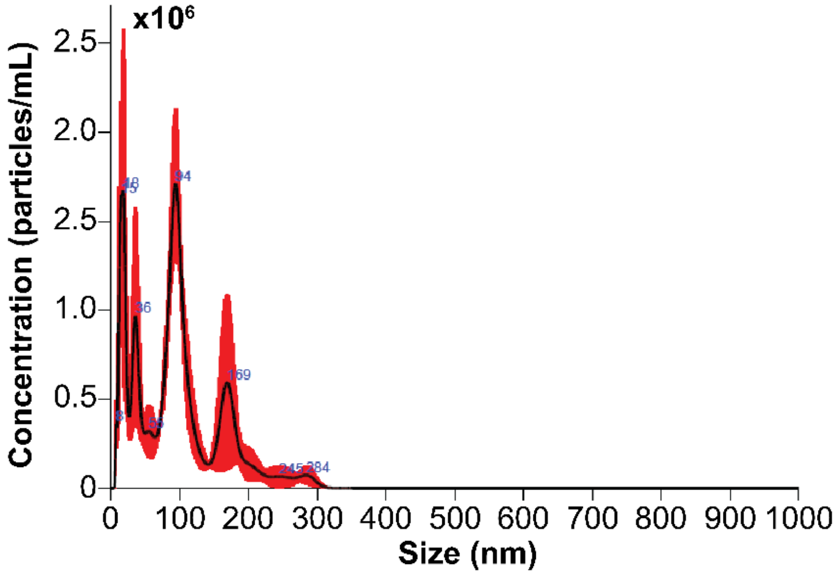


**Figure S1**. Nanoparticle tracking analysis of small extracellular vesicles (sEVs) purified from conditioned cell culture media using ultracentrifugation (UC).

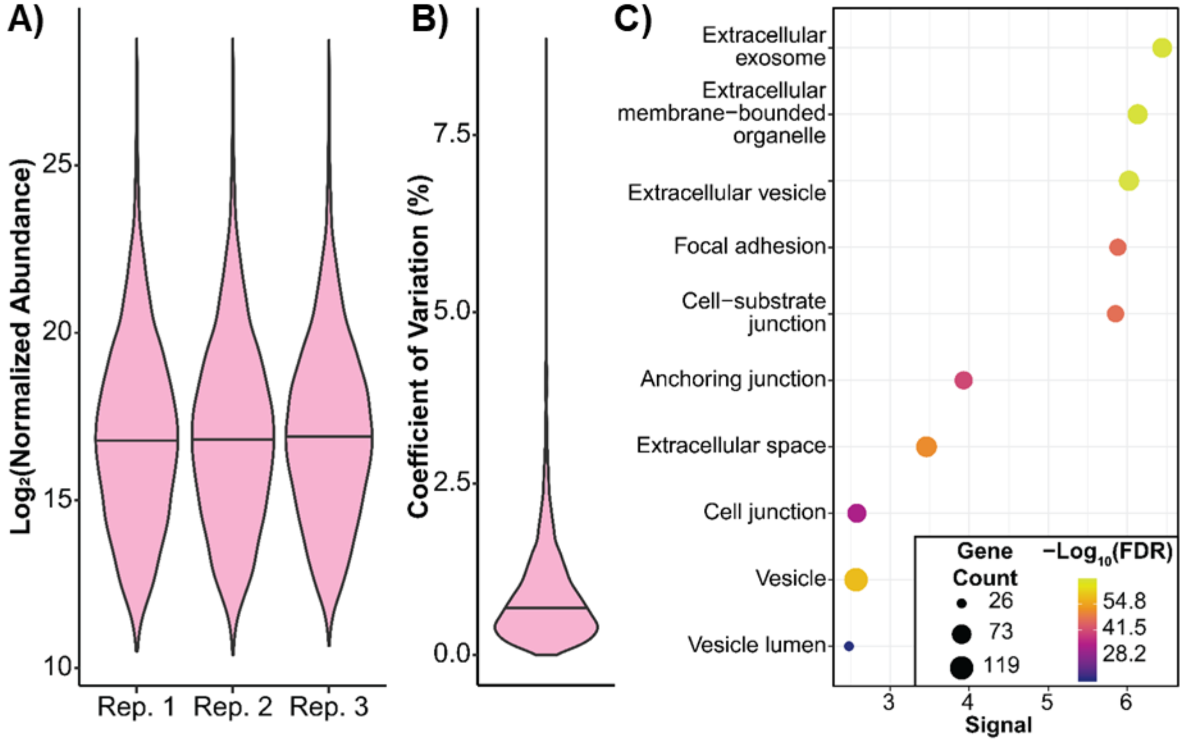


**Figure S2.** Proteomics analysis of small extracellular vesicles (sEVs) isolated from conditioned cell culture media. (**A**) Violin plots illustrate the distribution of values across the three replicates. (**B**) Violin plot demonstrates the distribution of coefficients of variation (%) within the dataset. (**C**) Gene Ontology Subcellular Localization (GOSCL) analysis of the top 200 most abundant proteins identified from cell culture sEVs.

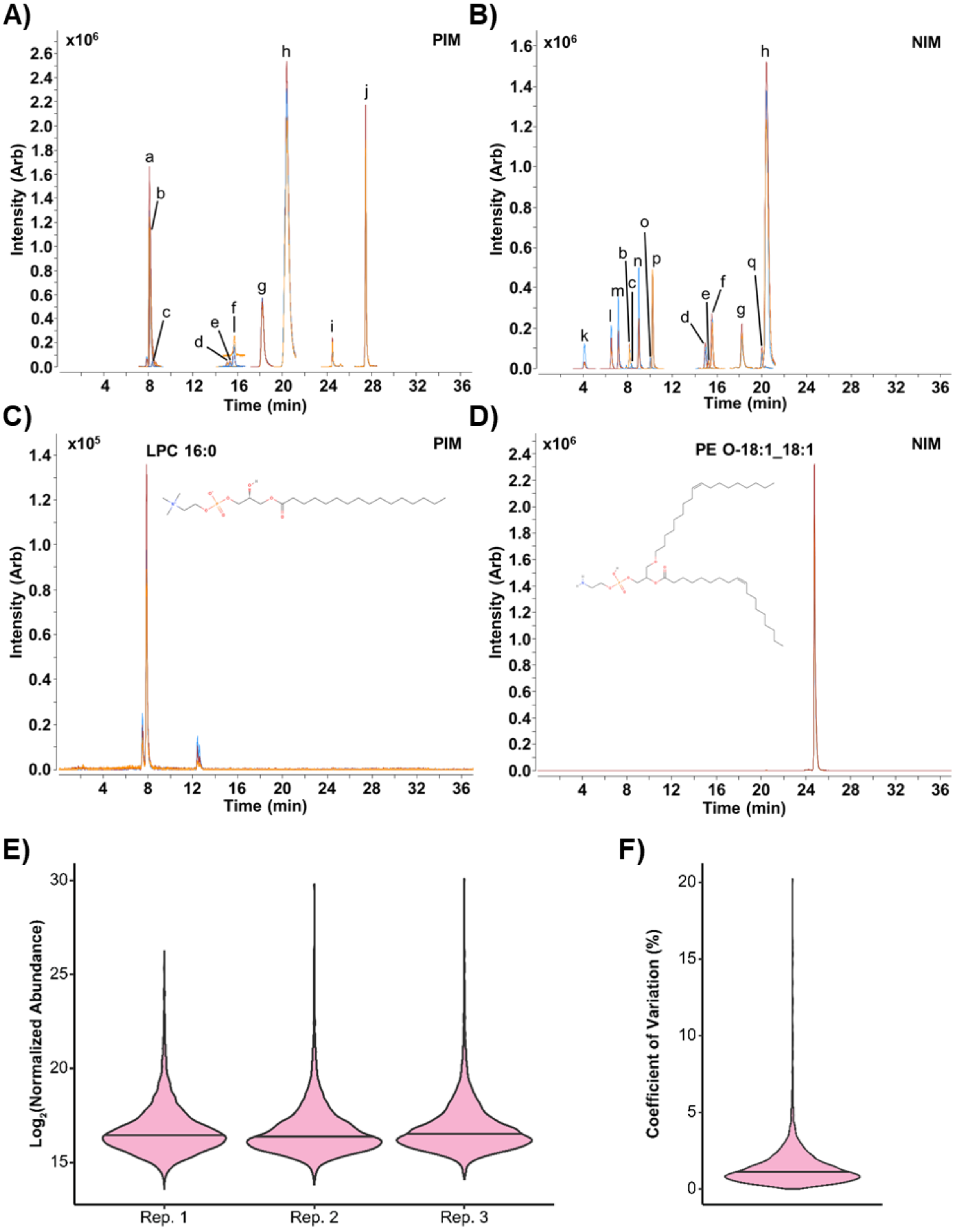


**Figure S3.** Lipidomics analysis of small extracellular vesicles purified from conditioned cell culture media. Overlay of extracted ion chromatograms (EICs) of isotopically labeled internal standards in (**A**) positive ion mode (PIM) and (**B**) negative ion mode (NIM). (**C**) EIC of lysophosphatidylcholine LPC 18:0 from PIM. (**D**) EIC of ether-type phosphatidylethanolamine PE O-18:1_18:1 from NIM. (**E**) Violin plots displaying the distribution of values across all three replicates. (**F**) Violin plot displaying the coefficient of variation (%) of the $Log_2$ transformed data across the dataset. **a**: oleoylcarnitine ($d_3$); **b**: lysophosphatidylcholine 18:1 ($d_7$); **c**: lysophosphatidylethanolamine 18:1 ($d_7$); **d**: phosphatidylinositol 15:0_18:1 ($d_7$); **e**: phosphatidylserine 15:0_18:1 ($d_7$); **f**: phosphatidylglycerol 15:0_18:1 ($d_7$); **g**: sphingomyelin d18:1_18:1 ($d_9$); **h**: phosphatidylcholine 15:0_18:1 ($d_7$); **i**: diacylglycerol 15:0_18:1 ($d_7$); **j**: triacylglycerol 15:0_18:1($d_7$)_15:0; **k**: fatty acid (FA) 12:0 ($d_{23}$); **l**: FA 14:0 ($d_{27}$); **m**: FA 16:1 ($d_{14}$); **n**: FA 18:1 ($d_{17}$); **o**: monoacylglycerol 18:1 ($d_7$); **p**: FA 18:0 ($d_{35}$); **q**: phosphatidylethanolamine 15:0_18:1 ($d_7$).

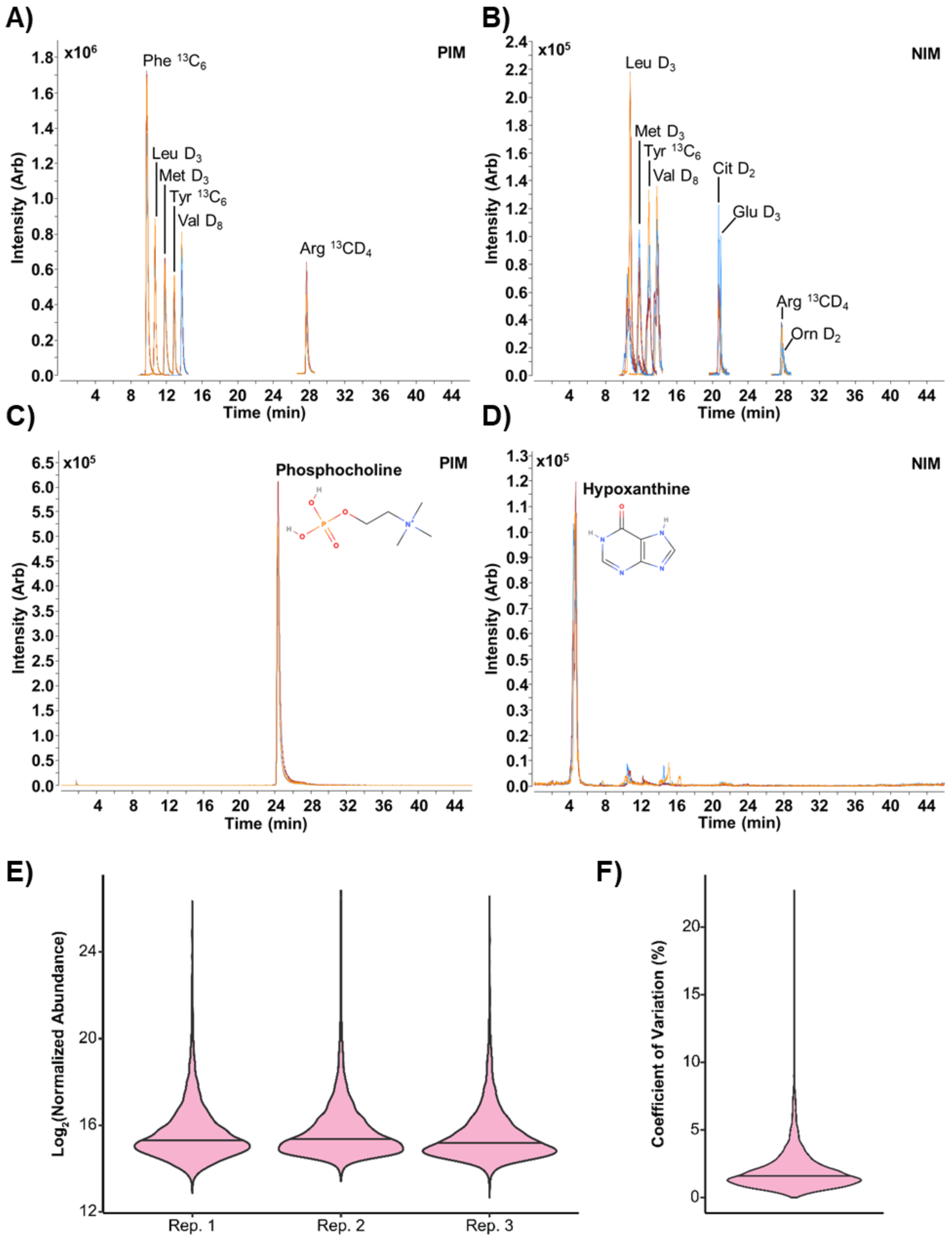


**Figure S4.** Metabolomics analysis of small extracellular vesicles (sEVs) isolated from conditioned cell culture media. (**A**) Overlay of extracted ion chromatograms (EICs) of isotopically labeled internal standards in (**A**) positive ion mode (PIM) and (**B**) negative ion mode (NIM). (**C**) EIC of phosphocholine from PIM. (**D**) EIC of hypoxanthine from NIM. (**E**) Violin plots demonstrate the distribution of values across the three replicates. (**F**) Violin plot demonstrates the distribution of the coefficients of variation (%) of the $Log_2$ transformed data within the dataset.

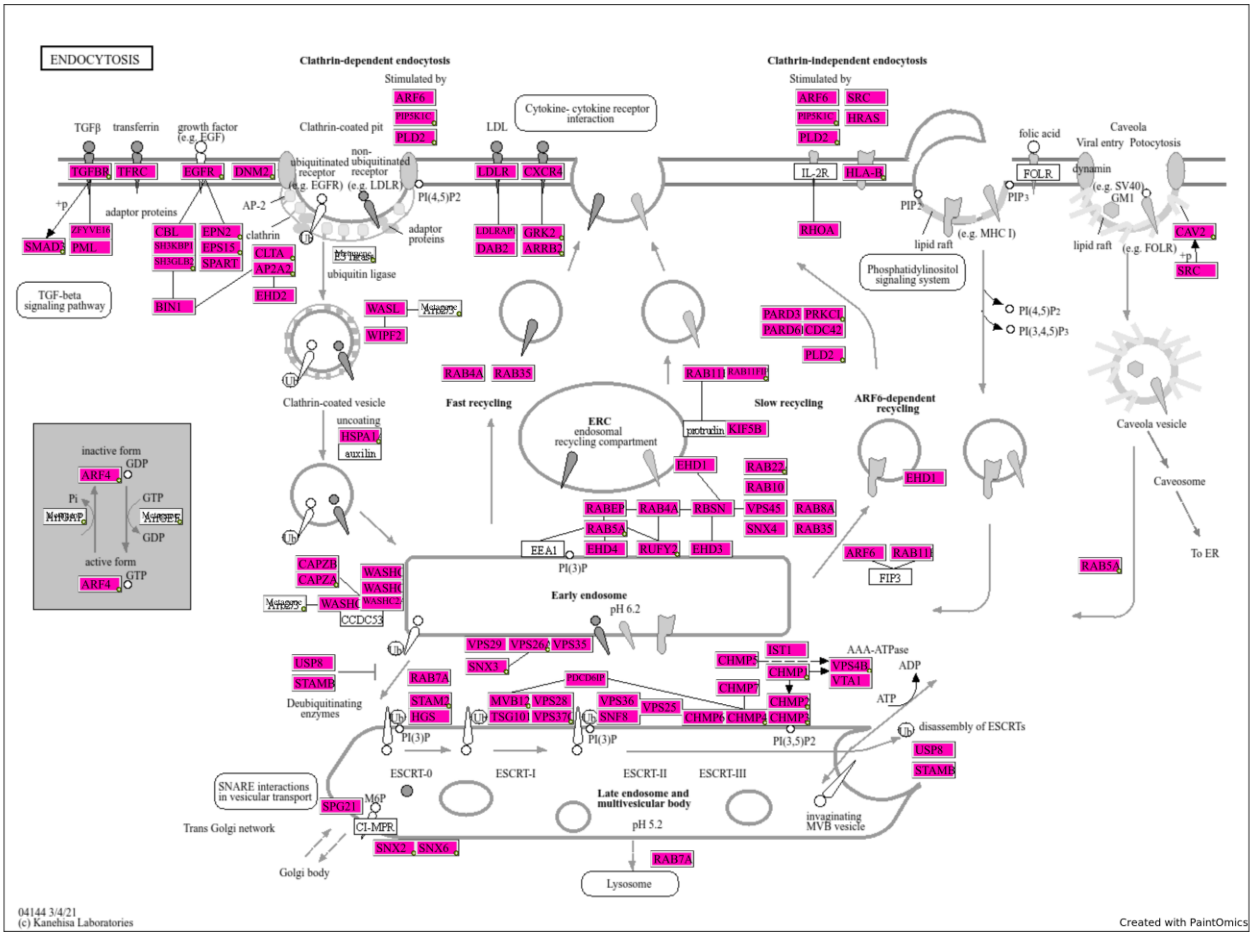


**Figure S5.** Coverage of molecules implicated in the endocytosis pathway. PaintOmics 4.0[9] was used to map which terms in the KEGG pathway,[10] “Endocytosis”, were identified in the multi-omics analysis of cell culture-derived sEVs. Terms highlighted in pink were detected in these analyses.

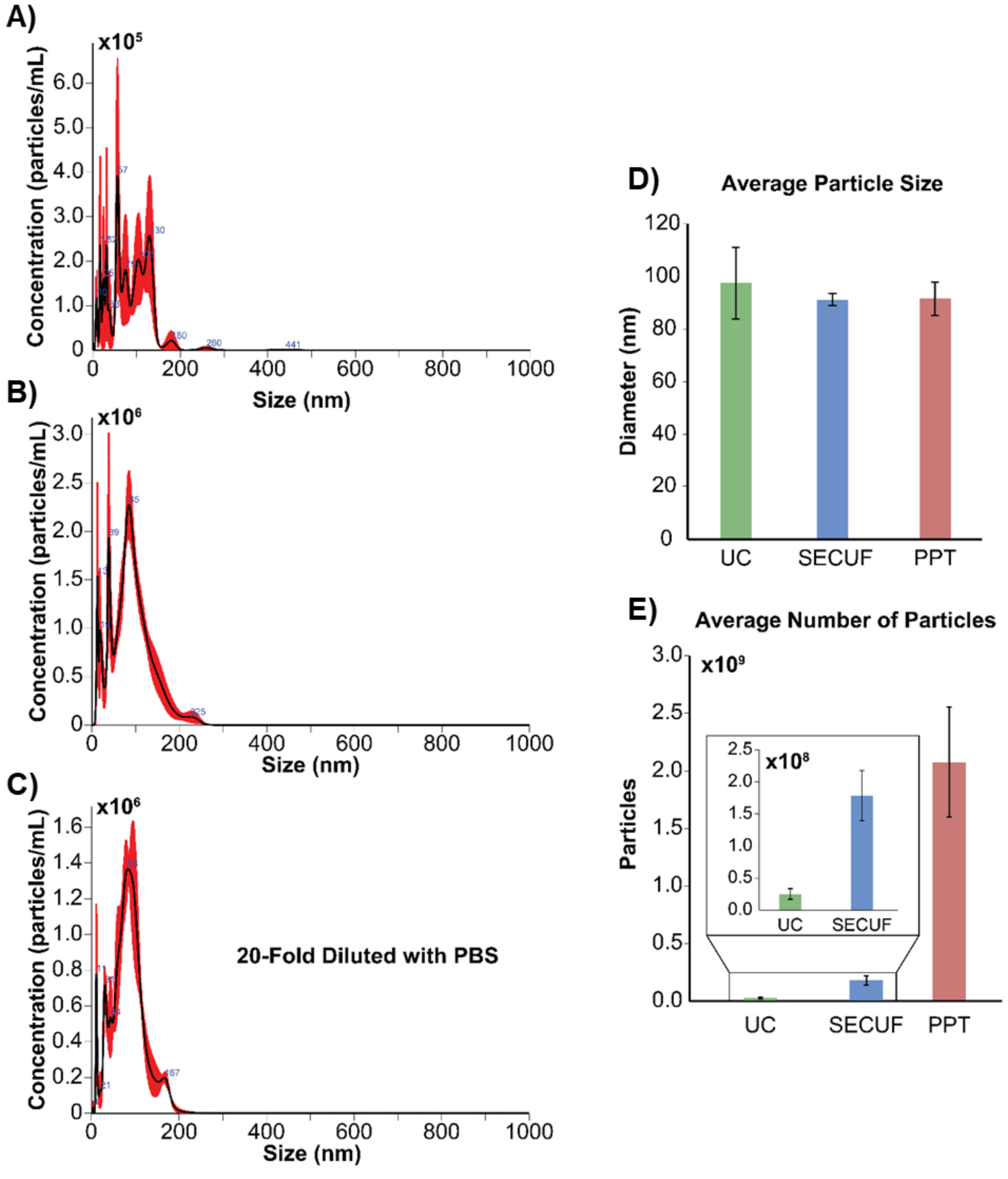


**Figure S6** Nanoparticle tracking analysis of small extracellular vesicles (sEVs) isolated from plasma using ultracentrifugation (UC), size exclusion chromatography with ultrafiltration (SECUF), and polymer precipitation (PPT). Representative plots displaying the size distribution of particles isolated using (**A**) UC, (**B**) SECUF, and (**C**) PPT. (**D**) Bar plots indicating the average particle sizes in populations purified using UC, SECUF, and

PPT. (**E**) Bar plots demonstrating differences in yield dependent on which purification approach was used.

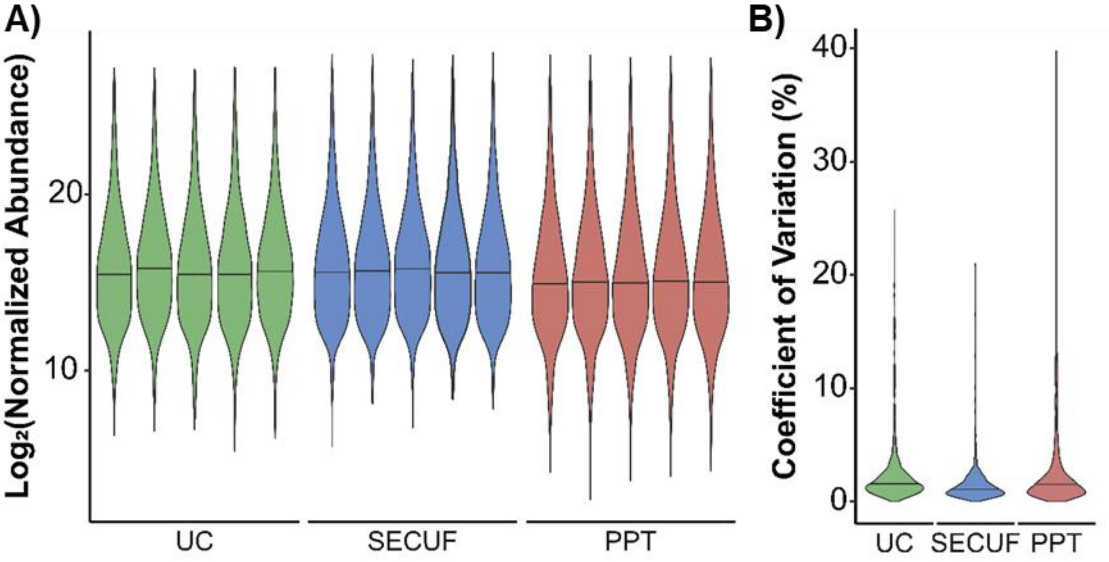


**Figure S7.** Proteomics of small extracellular vesicles (sEVs) purified from plasma using ultracentrifugation (UC), size exclusion chromatography with ultrafiltration (SECUF), and polymer precipitation (PPT). (**A**) Violin plots demonstrate the distribution of abundance values across samples from all conditions. (**B**) Distributions of coefficients of variation (%) of $Log_2$ transformed values in each of the three conditions.

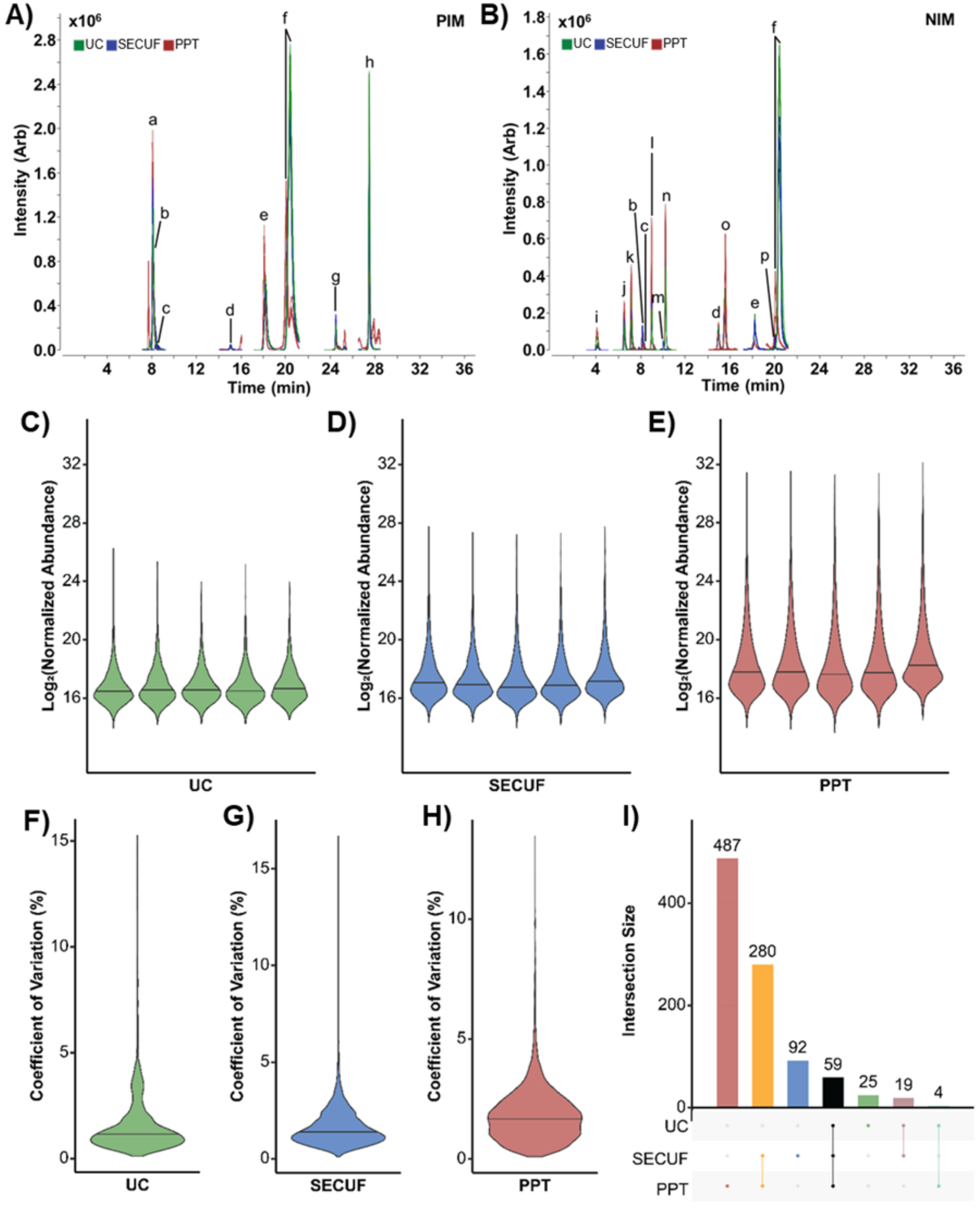


**Figure S8.** Lipidomics analysis of small extracellular vesicles purified from plasma using ultracentrifugation (UC), size exclusion chromatography with ultrafiltration (SECUF), and polymer precipitation (PPT). Overlays of extracted ion chromatograms of isotopically labeled internal standards detected in all samples in (**A**) positive ion mode and (**B**) negative ion mode. Violin plots illustrate the distribution of values in (**C**) UC samples, (**D**) SECUF samples, and (**E**) PPT samples. Violin plots illustrate the distribution of coefficients of variation (%) of the $Log_2$ transformed data for the (**F**) UC samples, (**G**) SECUF samples, and (**H**) PPT samples. (**I**) UpSet plot illustrates the degree of overlap in annotated features between the different conditions. **a**: oleoylcarnitine ($d_3$); **b**: lysophosphatidylcholine 18:1 ($d_7$); **c**: lysophosphatidylethanolamine 18:1 ($d_7$); **d**: phosphatidylinositol 15:0_18:1 ($d_7$); **e**: sphingomyelin d18:1_18:1 ($d_9$); **f**: phosphatidylcholine 15:0_18:1 ($d_7$); **g**: diacylglycerol 15:0_18:1 ($d_7$); **h**: triacylglycerol 15:0_18:1($d_7$)_15:0; **i**: fatty acid (FA) 12:0 ($d_{23}$); **j**: FA14:0 ($d_{27}$); **k**: FA 16:1 ($d_{14}$); **l**: FA 18:1 ($d_{17}$); **m**: monoacylglycerol 18:1 ($d_7$); **n**: FA 18:0 ($d_{35}$); **o**: phosphatidylglycerol 15:0_18:1 ($d_7$); **p**: phosphatidylethanolamine 15:0_18:1 ($d_7$).

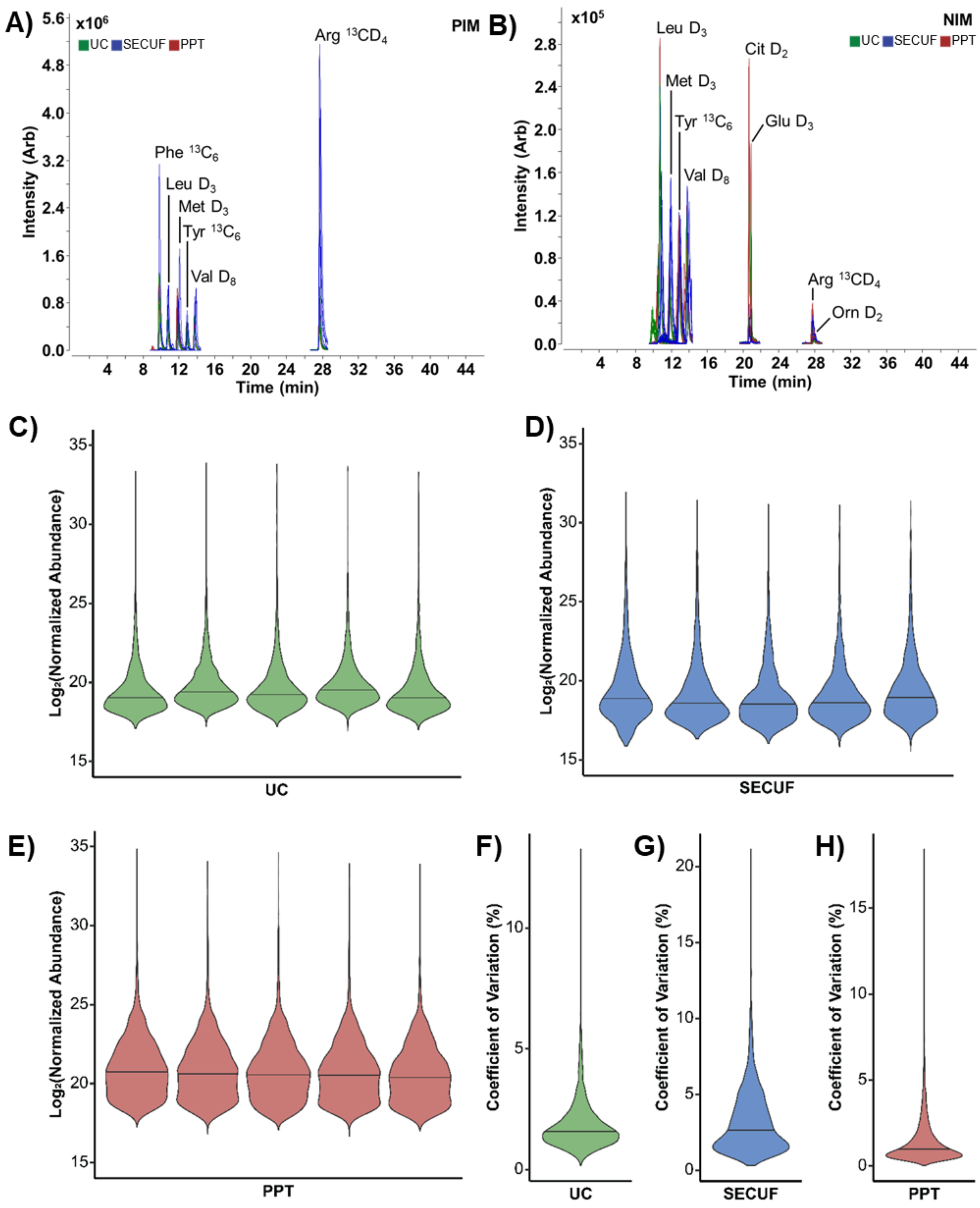


**Figure S9** Metabolomics analysis of small extracellular vesicles isolated from plasma using ultracentrifugation (UC), size exclusion chromatography with ultrafiltration (SECUF), and polymer precipitation (PPT). Overlays of extracted ion chromatograms of isotopically labeled internal standards in (**A**) positive ion mode and (**B**) negative ion mode.

Violin plots display the distribution of values across replicates in the (**C**) UC condition, (**D**) SECUF condition, and (**E**) PPT condition. Violin plots displaying the coefficient of variation (%) of $Log_2$ transformed values into the (**F**) UC dataset, (**G**) SECUF dataset, and (**H**) PPT dataset.

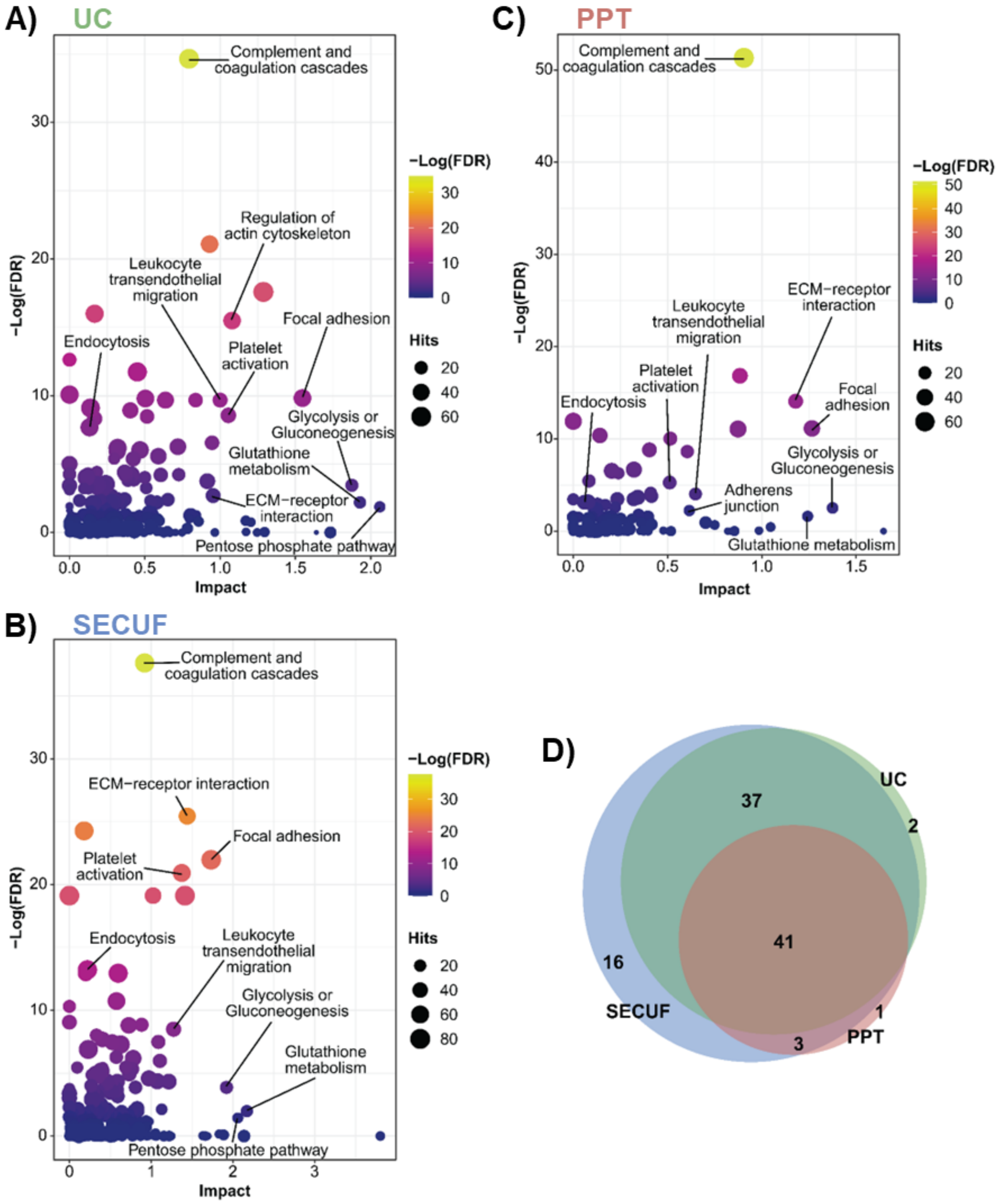


**Figure S10.** Joint pathway analysis of circulating small extracellular vesicles. (**A-C**) Joint pathway analysis performed using MetaboAnalyst[11] of identified lipids, metabolites, and proteins in small extracellular vesicles isolated from plasma using ultracentrifugation (UC), size exclusion chromatography with ultrafiltration (SECUF), and polymer precipitation (PPT), respectively. (**D**) A Venn diagram demonstrates overlap in significantly enriched (FDR < 0.05) pathways identified from UC, SECUF, and PPT purified sEVs.
